# Overexpressed *Malat1* Drives Metastasis through Inflammatory Reprogramming of Lung Adenocarcinoma Microenvironment

**DOI:** 10.1101/2023.03.20.533534

**Authors:** Elena Martínez-Terroba, Fernando J. de Miguel, Vincent Li, Camila Robles-Oteiza, Katerina Politi, Jesse R. Zamudio, Nadya Dimitrova

## Abstract

Metastasis is the main cause of cancer deaths but the molecular events leading to metastatic dissemination remain incompletely understood. Despite reports linking aberrant expression of long noncoding RNAs (lncRNAs) with increased metastatic incidence*, in vivo* evidence establishing driver roles for lncRNAs in metastatic progression is lacking. Here, we report that overexpression of the metastasis-associated lncRNA *Malat1* (metastasis-associated lung adenocarcinoma transcript 1) in the autochthonous K-ras/p53 mouse model of lung adenocarcinoma (LUAD) is sufficient to drive cancer progression and metastatic dissemination. We show that increased expression of endogenous *Malat1* RNA cooperates with p53 loss to promote widespread LUAD progression to a poorly differentiated, invasive, and metastatic disease. Mechanistically, we observe that *Malat1* overexpression leads to the inappropriate transcription and paracrine secretion of the inflammatory cytokine, Ccl2, to augment the mobility of tumor and stromal cells *in vitro* and to trigger inflammatory responses in the tumor microenvironment *in vivo*. Notably, Ccl2 blockade fully reverses cellular and organismal phenotypes of *Malat1* overexpression. We propose that *Malat1* overexpression in advanced tumors activates Ccl2 signaling to reprogram the tumor microenvironment to an inflammatory and pro-metastatic state.

## Introduction

Identification and functional characterization of long noncoding RNAs (lncRNAs) has suggested a potential wealth of unexplored cancer targets (Olivero and Dimitrova, 2020). While lncRNAs are rarely subject to recurrent mutations in cancer (Rheinbay et al., 2020), lncRNAs are frequently expressed at aberrant levels in tumors compared to normal tissues (Gutschner and Diederichs, 2012; Yan et al., 2015). Moreover, in many cases, the dysregulated expression of lncRNAs has been shown to strongly correlate with accelerated cancer progression, metastatic dissemination, and poor patient prognosis (Liu et al., 2021). Determining whether and how aberrantly expressed lncRNAs drive tumor progression has important clinical and therapeutic implications.

*Malat1* (metastasis-associated lung adenocarcinoma transcript 1) was first identified in a microarray screen of tumors from patients with non-small cell lung cancer and was found to be upregulated in the tumors with a higher propensity to metastasize (Ji et al., 2003; Muller-Tidow et al., 2004). Since these initial studies, *Malat1* overexpression has been reported in over 20 different solid and lymphoid tumors and consistently found to correlate with tumor progression and metastasis, highlighting its importance as a prognostic marker (Arun et al., 2020). In addition, genetic inactivation and transient knockdown of *Malat1* in autochthonous and xenograft murine models of breast and lung adenocarcinoma were found to significantly reduce metastatic dissemination (Arun et al., 2016; Gutschner et al., 2013). These studies provided evidence for a pro-metastatic function of *Malat1* and suggested downregulation of *Malat1* as a possible therapeutic strategy in metastatic disease.

Despite *Malat1* being a well-conserved, ubiquitously expressed, and highly abundant lncRNA (Hutchinson et al., 2007), its cellular functions and molecular mechanisms are poorly understood, which in turn has limited insights into its contribution to tumor development. For example, *Malat1* is a known component of alternative pre-mRNA splicing hubs called nuclear speckles and has been shown to associate with active chromatin (Hutchinson *et al*., 2007; Tripathi et al., 2010; West et al., 2014). Consistently, inactivation of *Malat1* in cellular and organismal models of cancer has been correlated with changes in the expression of genes linked to proliferation, differentiation, and cellular mobility (Arun *et al*., 2016; Gutschner *et al*., 2013). However, analyses of *Malat1*-deficient cells and tissues in three independent genetically engineered loss-of-function mouse models did not reveal any apparent defects in speckle formation and did not uncover any significant changes in alternative splicing or gene expression patterns (Eissmann et al., 2012; Nakagawa et al., 2012; Zhang et al., 2012).

To reconcile the cancer roles of *Malat1* with its apparent redundancy under normal physiological conditions, it has been proposed that *Malat1* may act in a context-dependent manner (Arun *et al*., 2020; Sun and Ma, 2019). However, gain-of-function approaches aimed at recapitulating the consequences of *Malat1* overexpression in the context of cancer progression have been limited and have led to conflicting results. On the one hand, overexpression of a *Malat1* fragment was found to be sufficient to increase anchorage independent growth of primary mouse embryonic fibroblasts and to induce metastatic dissemination of a non-metastatic murine mammary cancer cell line (Gao et al., 2014; Li et al., 2009). On the other hand, whole-body transgenic overexpression of *Malat1* in an autochthonous murine model of metastatic breast cancer unexpectedly led to reduced incidence of lung metastases (Kim et al., 2018).

To overcome limitations of previous approaches and to directly address the contribution of *Malat1* overexpression to tumorigenesis, in this study we describe the development and characterization of a novel CRISPR-based approach to overexpress endogenous *Malat1* in autochthonous LUAD mouse models and in patient derived LUAD cell lines. Our findings conclusively establish that overexpressed *Malat1* is a powerful driver of tumor progression and reveal an unanticipated paracrine mechanism by which *Malat1*-overexpressing tumor cells signal to innate immune cells to reprogram the tumor microenvironment to an inflammatory and pro-metastatic state. This study brings to light a central role for a dysregulated lncRNA in driving metastatic progression, and uncovers a therapeutically accessible target, Ccl2, as the central mediator of metastasis downstream *Malat1* overexpression.

## Results

### Overexpressed *Malat1* is a conserved feature of human and murine LUAD

As the predominant form of processed *Malat1* is non-polyadenylated (Wilusz et al., 2008), accurate quantification of *Malat1* levels in polyA-selected tumor sequencing datasets is challenging (Fig. S1A). To independently evaluate the reported link between elevated *Malat1* levels in lung adenocarcinoma (LUAD) and disease progression, we used RNAScope to quantify the abundance of processed *Malat1* in lung cancer samples on a tissue microarray (TMA) (Fig. S1B, C and Table S1). We confirmed increased levels of *Malat1* in tumors compared to adjacent normal tissue (Fig. 1A, B) and observed that high *Malat1* is strongly predictive of decreased disease-free survival (p=0.012, Fig. 1C and Tables S2 and S3), consistent with previous studies and supporting a role in tumor development (Mei et al., 2019).

**Figure 1.**
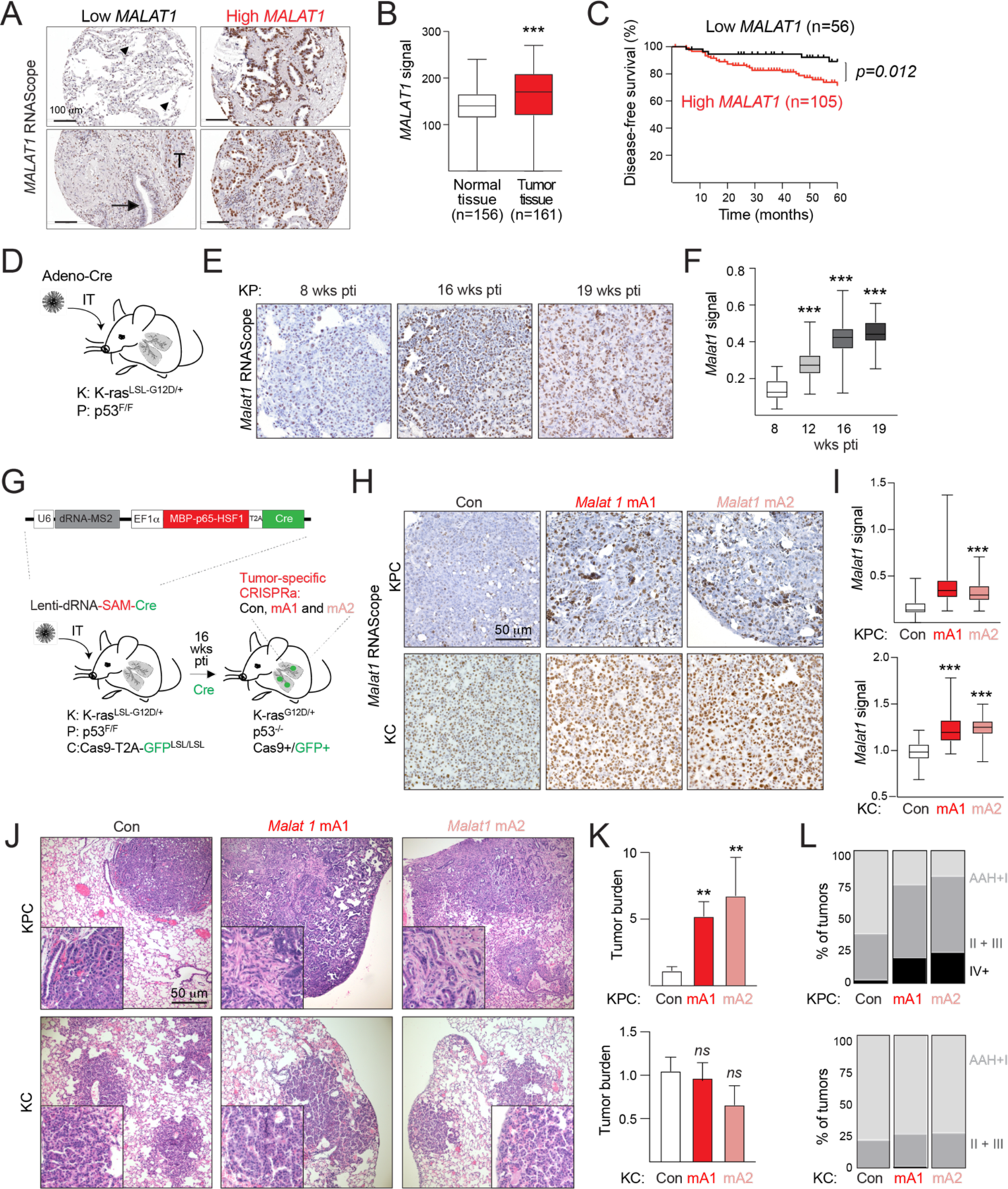
*Malat1* overexpression in autochthonous KP LUAD model cooperates with p53 to drive tumor progression. (A-C). High expression of *Malat1* is predictive of poor prognosis in lung cancer patients. (A) Representative images of RNAScope detection of human *Malat1* RNA in normal and tumor tissue from lung cancer patients tissue microarrays; arrowheads, alveoli; arrow bronchiole; T, tumor; (B) Boxplot of human *Malat1* RNA levels in normal and tumor tissue (n>150 tumors, unpaired t-test, p < 0.001); (C) Kaplan–Meier survival curves. Patients were stratified in two groups according to *Malat1* signal (Low *Malat1*, bottom tertile, n=56, High *Malat1*, top two tertiles, n=105; log-rank test, p=0.012); (D) Schematic of KP model of LUAD, highlighting tumor initiation via intratracheal infection with adenoviral Cre recombinase (Adeno-Cre); (E) Representative images of murine *Malat1* levels determined by RNAScope in tumors from mice in (D) sacrificed at indicated time-points post tumor initiation (pti); (F) Quantification using QuPath software of murine *Malat1* signal in tumors from images in (E) (8 wks pti, n=15; 12 wks pti, n=33; 16 wks pti, n=52; 19 wks pti, n=47 tumors); (G) Schematic of tumor-specific *Malat1* CRISPRa in KC and KPC mouse models of LUAD; (H-I) RNAScope analysis (H) and quantification of murine *Malat1* RNA levels in KC and KPC tumors (n>50 tumors, unpaired t-test) (I); (J) H&E staining of lung sections of KC and KPC mice infected with indicated dRNAs and analyzed at 16 weeks pti; (K) Quantifications of relative tumor burden of mice in (J) (KPC: Con, n=8; mA1, n=9; mA2, n=8; KC: Con, n=10; mA1, n=12; mA2, n=8; unpaired t-test); (L) Quantification of tumor grade in mice in (J) (n>150 tumors).

The *K-ras^LSL-G12D/+^; p53^F/F^* (KP) mouse is a powerful genetic tool to study LUAD initiation and progression (Jackson et al., 2005; Jackson et al., 2001; Winslow et al., 2011). In the KP model, Cre-mediated activation of the G12D oncogenic mutation in Kras and deletion of the p53 tumor suppressor induces synchronous development of LUAD lesions in the normal lung microenvironment (DuPage et al., 2009). Interestingly, RNAScope quantification of murine *Malat1* in KP tumors revealed a progressive increase of *Malat1* levels over the course of tumor development (Fig. 1D-F). The consistent increase of *Malat1* levels in human and mouse LUAD suggested that overexpressed *Malat1* may play a conserved role in LUAD progression.

### *Malat1* overexpression is sufficient to drive LUAD progression and metastasis

To directly test the functional role of elevated *Malat1* expression in LUAD at the organismal level, we modelled *Malat1* overexpression in KP mice by performing tumor-specific overexpression of endogenous *Malat1* using CRISPR activation (CRISPRa) at the time of tumor initiation. To this end, we crossed K and KP mice to the *Rosa26-Cas9-T2A-GFP^LSL/LSL^* (C) allele (Platt et al., 2014) to generate KC and KPC cohorts enabling Cre-mediated tumor-specific expression of Cas9 (Fig. 1G). We also constructed a trifunctional lentiviral vector to express (1) U6-driven control (Con) or two independent *Malat1*-specific (mA1 and mA2) dead RNAs extended with MS2 repeats (dRNAs), (2) the SAM (Synergistic Activation Meditator) components fused to the MS2 binding protein (MBP) (Dahlman et al., 2015), and (3) Cre recombinase, required for tumor initiation (Lenti-dRNA-SAM-Cre; Fig. 1G). At 16 weeks post tumor initiation, we confirmed by RNAScope significant tumor-specific upregulation of *Malat1* in KPC and KC animals with the two independent *Malat1*-targeting dRNAs compared to a control dRNA (Fig. 1H, I).

To evaluate the consequences of *Malat1* overexpression, we performed histopathological analysis of tumor-bearing lungs from *Malat1*-overexpressing and control KPC and KC mice sacrificed at 16 weeks post tumor initiation. Quantification of H&E images revealed that *Malat1* overexpression with the two independent dRNAs led to a 5-6-fold increase of tumor burden in KPC mice (Fig. 1J, K). In addition, while lungs in control KPC animals presented with primarily atypical adenomatous hyperplasia (AAH) and low grades (I, II, and III) tumors (DuPage *et al*., 2009), over 20% of *Malat1*-overexpressing tumors progressed to high grade (IV+) (Fig. 1L). Notably, high grade tumors in *Malat1*-overexpressing KPC animals were characterized by glandular architecture, extensive desmoplasia, and evidence of local invasion but did not correspond to increased cellular proliferation (Fig. S2A-C). These data demonstrated that overexpressed *Malat1* does not significantly impact tumor growth but is a potent driver of tumor progression. In contrast, *Malat1* overexpression did not significantly affect overall tumor burden or grade in KC animals (Fig. 1J-L), indicating that p53 function is a barrier to *Malat1*-driven tumor progression.

### Changes of tumor microenvironment and tumor cells accompany LUAD progression and metastatic dissemination

Strikingly*, Malat1*-overexpressing KPC animals had a significantly increased incidence of metastases. At 16 weeks post tumor initiation, over 50% of mA1-and mA2-expressing animals presented with local metastases in the thoracic cavity and local lymph nodes and over 25% showed evidence of distal metastatic dissemination to the liver and distal lymph nodes (Fig. 2A, B). At the same time-point, fewer than 25% of control animals presented with only local metastases, consistent with previous data in this model (Fig. 2B) (DuPage *et al*., 2009; Winslow *et al*., 2011). Consequently, *Malat1* overexpression was associated with significantly shorter overall survival compared to control animals (Fig. 2C; mA1, p=0.0283; mA2, p=0.0198). The accelerated tumor progression of *Malat1*-overexpressing KPC tumors correlated with significant changes in previously established markers of metastatic progression, namely Nkx2.1 (also known as Ttf1) loss and Hmga2 overexpression (Fig. 2D-F) (Winslow *et al*., 2011). As above, KC tumors with a functional p53 pathway did not present Hmga2 overexpression (Fig. 2G, H), supporting a key role for p53 in restraining tumor progression resulting from *Malat1* overexpression. We concluded that *Malat1* overexpression in murine KP LUAD cooperates with p53 loss to drive aggressive disease and metastasis, resulting in phenotypes consistent with LUAD patient disease progression.

**Figure 2.**
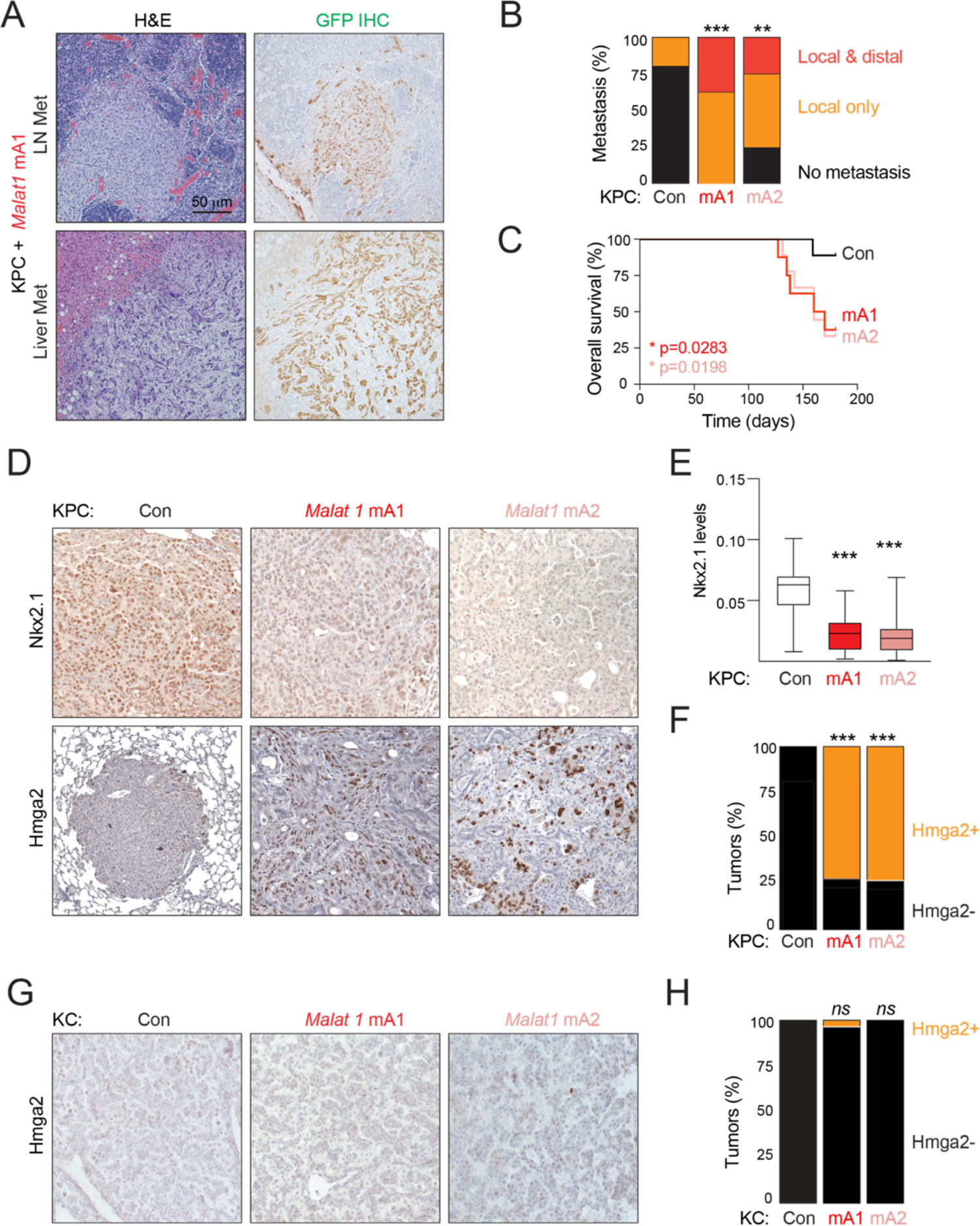
*Malat1* overexpression enables LUAD metastatic dissemination. (A) Representative H&E and GFP IHC staining of liver and lymph node (LN) metastasis (Met) of the long-term experiment of KPC animals infected with mA1 dRNA; (B) Quantification of metastasis of the mice analyzed in (A) (n=10 mice per group, unpaired t test); (C) Survival curves of the animals shown in (A). Log-rank test was used for statistical analysis. Presence of ethical endpoint criteria were used as endpoint; (D) IHC staining for the metastasis markers Nkx2.1 and Hmga2 of KPC animals analyzed in (A); (E) Quantification of Nkx2.1 staining in images in (D) (n>20 tumors, unpaired t-test); (F) Contingency analysis showing the percentage of positive tumors for Hmga2 immunostaining (Chi-square 28.68, P<0.001, n=79 tumors); (G) IHC staining for Hmga2 marker in the KC the cohort; (H) Contingency analysis showing the percentage of positive tumors for Hmga2 immunostaining in KC cohort (Fisher exact test P=0.504, n=157 tumors).

### *Malat1* overexpression downregulates epithelial transcription programs

Previous studies have proposed that *Malat1* contributes to tumor progression through the activation of epithelial mesenchymal transition (EMT) transcription programs (Arun *et al*., 2016; Guo et al., 2015; Gutschner *et al*., 2013; Tano et al., 2010). To investigate whether *Malat1* overexpression drives metastatic progression through an EMT mechanism in LUAD, we performed RNA sequencing of Live/GFP+/CD45-/Epcam+ cancer cells isolated from dissociated Con- and mA1-expressing KPC tumors at 16 weeks post tumor initiation (Fig. 3A). Principal component analysis (PCA) revealed that the first component (PC1), which accounts for 34% of variation between the samples, correlated with *Malat1* overexpression (Fig. 3A). Gene set enrichment analysis (GSEA) of PC1 showed downregulation of Cdh1 (also known as E-cadherin) target genes (ONDER_CDH1_TARGETS_2_DN), as well as enrichment of metastatic programs (JAEGER_METASTASIS_DN, WU_CELL_MIGRATION, RICKMAN_METASTASIS_DN, Fig. 3B and Table S4). Next, analysis of differentially expressed genes (DEGs) (|log2FC|>1; p<0.05) revealed that 227 genes were consistently downregulated and 98 genes were consistently upregulated in mA1 compared to Con KPC tumors, suggesting that *Malat1* overexpression alters the cancer transcriptome (Fig. 3C and Tables S5 and S6). GSEA of downregulated genes pointed to a strong enrichment of genes involved in epithelial cell differentiation and epithelial development (GO_CELL_ADHESION, GO_EPITHELIAL_CELL_DIFFERENTIATION, GO_EPITHELIUM_DEVELOPMENT) (Fig. 3D). Consistently, cumulative frequency plots confirmed downregulation of genes involved in epithelial differentiation (p=0.000164, GO_EPITHELIAL_CELL_DIFFERENTIATION) and suppressors of metastasis (p=1.1e-7, CGP_RICKMAN_METASTASIS_DN, Fig. 3E) and GSEA supported a significant enrichment of EMT genes (p=0.0221, HALLMARK_EPITHELIAL_ MESENCHYMAL_TRANSITION, Fig. 3F and Table S7). The downregulation of epithelial gene expression programs was due to changes at the transcriptional level, rather than due to post-transcriptional processes, as we observed comparable genesets enriched in analyses of both exonic or intronic read coverage (Fig. 3G, H and Table S8). Of note, transcriptomics data did not lend support to several previously proposed models for *Malat1* activities, including miRNA-mediated mechanisms, indirect effects through deregulation of neighboring genes (Fig. S3), or splicing control (Arun *et al*., 2020; Sun and Ma, 2019). These results supported previously proposed models that linked *Malat1* and EMT but clarified that *Malat1* overexpression primarily leads to the transcriptional downregulation of epithelial transcription programs.

**Figure 3.**
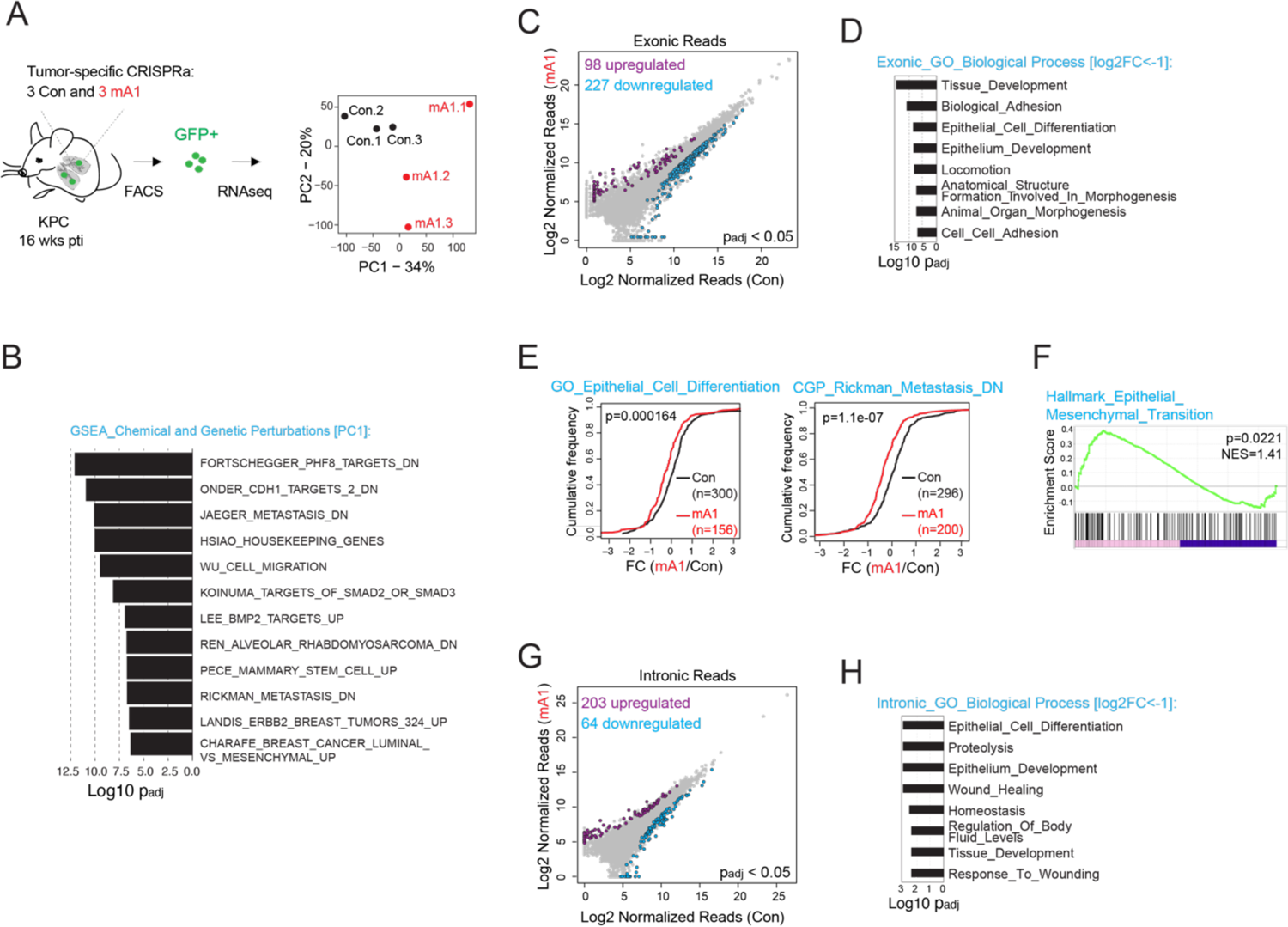
*Malat1* overexpression downregulates epithelial gene expression programs in LUAD. (A) Schematic of RNA isolation from GFP+/Cd45-/EpCam+/Live Con and mA1 KPC tumor cells and PCA of RNA sequencing data; (B) Enrichment of GSEA Chemical and Genetic Perturbations genesets in PC1 genes; -log10(padj) on x-axis; (C) Scatter plot of DEGs determined from exonic read coverage (purple, upregulated, blue, downregulated, |log2FC|>1, adjusted p<0.05); (D) Enrichment of Gene Ontology (GO) Biological Process genesets in downregulated genes in (C); (E) Cumulative frequency distribution plots show differential regulation of indicated genesets relative to matched control genesets in mA1 compared to Con samples; (F) GSEA of genes differentially expressed in mA1 compared to Con samples showing enrichment of the Hallmark Epithelial Mesenchymal Transition geneset; (G) Scatter plot of DEGs determined from intronic read coverage in RNA sequencing of Con and mA1 KPC tumors (purple, upregulated, blue, downregulated, |log2FC|>1, adjusted p<0.05); (H) Enrichment of Gene Ontology (GO) Biological Process genesets downregulated in mA1 compared to Con samples (log2FC<-1), showing similar pathways compared to the genesets obtained from exonic reads in (D).

### Paracrine effects of *Malat1* overexpression

Metastatic progression results from cumulative changes in both tumor cells and tumor stroma. Given the striking increase in frequency of metastatic dissemination in *Malat1*-overexpressing mice, we considered whether overexpressed *Malat1* may also contribute to the reprogramming of the tumor microenvironment to a pro-metastatic state. Consistent with this model, we observed that in addition to downregulation of epithelial signatures in tumor cells, illustrated by a significant decrease of the epithelial marker Cdh1 (Fig. 4A, B), *Malat1*-overexpressing tumors exhibited notable changes in the tumor stroma (Fig. 4A). We observed a 5-fold increase in the recruitment of α-Sma-positive cancer associated fibroblasts (CAFs) and increased abundance of Mason’s Trichrome-positive extracellular matrix (ECM) in *Malat1*-overexpressing tumors compared to controls (Fig. 4A, C). As CAF recruitment and ECM accumulation are associated with advanced LUAD grade, we concluded that *Malat1* overexpression correlates with pro-metastatic changes in the tumor microenvironment.

**Figure 4.**
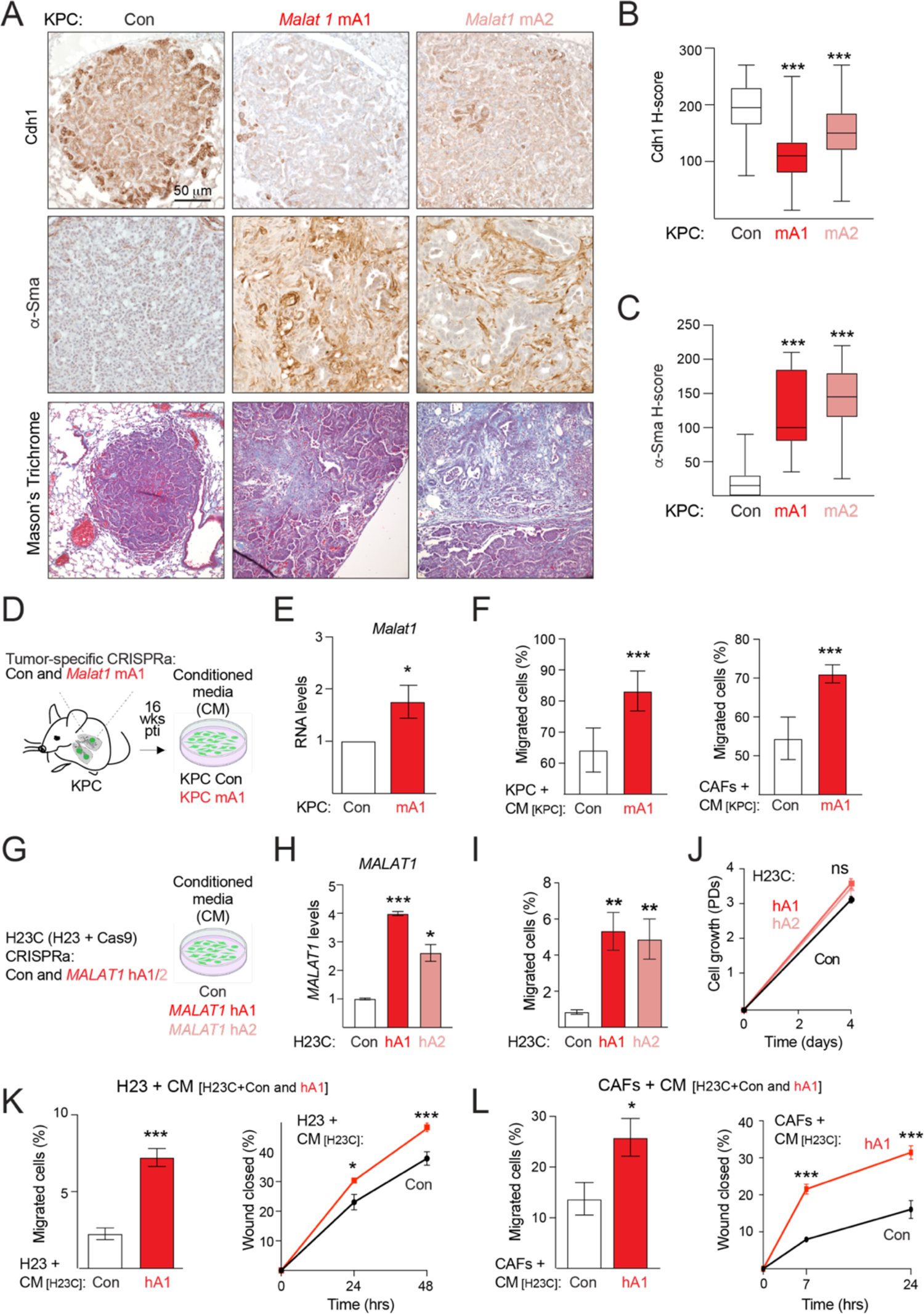
*Malat1* overexpression reprograms tumor stroma and stromal cell behavior through a paracrine mechanism. (A) Immunostaining of Cdh1, α-Sma, and Masson’s Trichrome staining for stroma infiltration in KPC animals infected with mA1 or mA2 dRNA; (B) Quantification of Cdh1 staining (n>20 tumors, unpaired t-test); (C) Quantification of α-Sma staining (n>20 tumors, unpaired t-test); (D) Schematic of the isolation of Con and mA1 KPC cell lines; (E) *Malat1* and Cdh1 RNA levels in indicated KPC cells; (F) Quantification of transwell migration by Boyden chamber assay of KPC cells or CAFs incubated with CM from KPC cells; (G) CRISPRa of human *Malat1* in H23 cells, transduced with Cas9 (H23C); (H) *Malat1* and CDH1 RNA levels detected in indicated cells; (I) Quantification of transwell migration by Boyden chamber assay of indicated H23C cells; (J) Growth analysis of cells in (I); (K) Quantification of cell migration by Boyden chamber (*left*) and wound healing (*right*) assays of H23C cells incubated with CM from H23C+Con or H23C+hA1 cells; (L) Quantification of cell migration by Boyden chamber (*left*) and wound healing (*right*) assays of CAFs incubated with CM from indicated cells. Data in this figure represented as mean ± SEM of biological replicates, n≥3, paired t-test.

We considered whether paracrine effects of *Malat1*-overexpressing cells may promote the observed pleiotropic changes in the tumor stroma. To first test this hypothesis *in vitro*, we established murine cell lines from KPC Con and mA1 tumor-bearing lungs and confirmed by quantitative PCR *Malat1* overexpression in mA1 compared to Con KPC cells (Fig. 4D, E). We next collected conditioned media (CM) from these cells and observed that CM from *Malat1-* overexpressing cells increased the transwell migration of both tumor cells and CAFs compared to CM collected from control KPC cells (Fig. 4F). These results revealed that overexpression of *Malat1* in our model indeed exerts paracrine effects on the behavior of neighboring cells, including stromal cells.

We also examined whether overexpression of human *Malat1* in patient-derived LUAD cell lines exerted similar effects. We performed CRISPRa of *Malat1* in the non-metastatic cell lines H23C (H23 expressing Cas9) and H2122C (H2122 expressing Cas9) with two independent dRNAs targeting human *Malat1*, hA1 and hA2 (Fig. 4G). We observed 2-9-fold increase in *Malat1* levels in hA1- and hA2-expressing cells compared to controls (Fig. 4H and S4A). *Malat1* overexpression did not affect cellular proliferation but was accompanied by a 5-fold increase in tumor cell mobility (Fig. 4H-J and S4B-D). Conversely, 50% downregulation of *Malat1* by CRISPR inhibition (CRISPRi) in the metastatic cell line H2009C (H2009 expressing Cas9) with two independent *Malat1*-targeting dRNAs, hI1 and hI2, led to a 2-3-fold decrease in tumor cell mobility compared to a control dRNA without impacting growth rate (Fig. S4E-H). Importantly, we observed that, analogous to murine LUAD cells, CM collected from *Malat1*-overexpressing but not control H23C cells was sufficient to stimulate transwell migration and accelerated wound healing of both H23 cells (Fig. 4K) and CAFs (Fig. 4L). Together, these findings revealed that *Malat1* overexpression in both murine and human LUAD cells promotes the mobility of tumor cells and CAFs through a soluble factor.

### RNA-based mechanism of overexpressed *Malat1*

LncRNAs can exert their activities either through the act of transcription from the lncRNA locus or through the accumulation of functional RNA molecules (Winkler and Dimitrova, 2021). Since CRISPRa increases both the transcription of *Malat1* and the mature RNA levels, we sought to distinguish between the two models by targeting the stability of *Malat1* post-transcriptionally. *Malat1* is transcribed as a polyadenylated transcript but cleaved by RNAse P to release a 5’ fragment, which is stabilized by a triple-helical terminal expression and nuclear retention element (ENE) (Fig. S1A) (Brown et al., 2014; Wilusz *et al*., 2008; Wilusz et al., 2012). To examine the importance of mature *Malat1* accumulation, we used two independent approaches to disrupt ENE folding. On one hand, we designed a guide RNA (gRNA) to recruit Cas9 to *Malat1* ENE genomic sequence and introduce CRISPR indels that might alter pairing interactions in the ENE (Fig. 5A, B). On the other hand, we designed a non-degrading antisense oligonucleotide (ASO) complementary to the *Malat1* ENE that would interfere with the folding of the protective triple-helical structure (Fig. 5C). Introduction of the ENE-targeting gRNA and ASO in two independent H23C cell lines overexpressing *Malat1* (H23C+hA1 and H23C+hA2) led to a 40-60% reduction of *Malat1* levels relative to corresponding non-targeting gRNA and ASO controls, confirming the stabilizing role of the ENE (Fig. 5D, E). Importantly, transwell migration assays revealed that both ENE mutagenesis and ENE interference significantly reduced the mobility of *Malat1*-overexpressing cells (Fig. 5F, G). These results indicated that the effects of *Malat1* overexpression can be reversed by reducing *Malat1* levels and therefore were due to increased *Malat1* RNA levels rather than resulting from altered transcription of *Malat1*.

**Figure 5.**
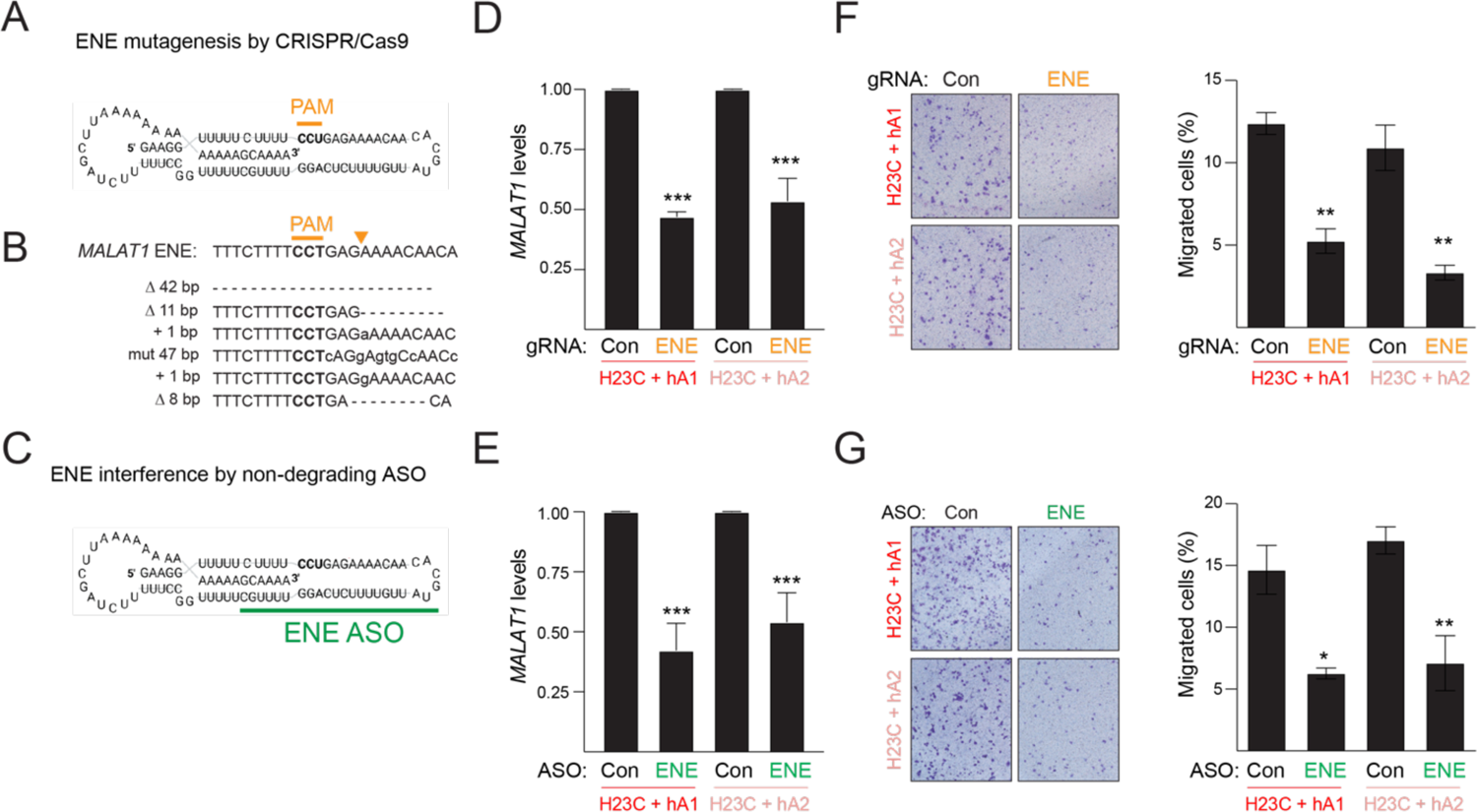
Increased accumulation, rather than production, of *Malat1* RNA mediates phenotypes of *Malat1* overexpression. (A) Schematic of the sequence and structure of *Malat1* ENE domain indicating the PAM site for CRISPR mutagenesis (yellow); (B) Analysis of ENE mutagenesis. *Top*, schematic indicating the PAM site (bold) and the Cas9 cleavage site (yellow arrow). *Bottom*, mutant alleles, determined by Sanger sequencing of H23C+hA1 cells expressing the ENE-targeting gRNA; (C) Schematic of the sequence and the non-degrading ASO targeting sequence (green) of *Malat1* ENE domain; (D-E) RT-qPCR analysis of *Malat1* levels in H23C hA1 and H23C hA2, expressing ENE-targeting gRNA (Mut) (D) or ENE-targeting ASO (E) relative to corresponding control non-targeting gRNA or ASO (Con); (F) Representative images and quantification of transwell migration by Boyden chamber assay of H23C+hA1 or H23+hA2 cells expressing ENE-targeting gRNA (ENE) compared to control gRNA (Con), (G) Representative images and quantification of transwell migration by Boyden chamber assay of H23C+hA1 or H23+hA2 cells treated with ENE-targeting ASO (ENE) compared to non-targeting ASO (Con). Data show mean ± SEM (n = 3, biological replicates).

### Secretion of inflammatory cytokine Ccl2 mediates the increased mobility of *Malat1* **overexpressing cells**

We next sought to identify the soluble factor(s) that mediate the effects of *Malat1* overexpression by profiling the proteome of CM collected from Con and hA1 H23C cells using antibody arrays (Fig. S5). We determined that the cytokine Ccl2, also known as Mcp1 (monocyte chemoattractant protein 1), a known tumor microenvironment modulator and metastasis enabler (Saji et al., 2001; Ueno et al., 2000; Valkovic et al., 1998), is enriched over 4-fold in CM from H23C cells overexpressing *Malat1* compared to controls (Fig. 6A, B) and correlated with a 10-15-fold upregulation of *Ccl2* RNA in these cells (Fig. 6C).

**Figure 6.**
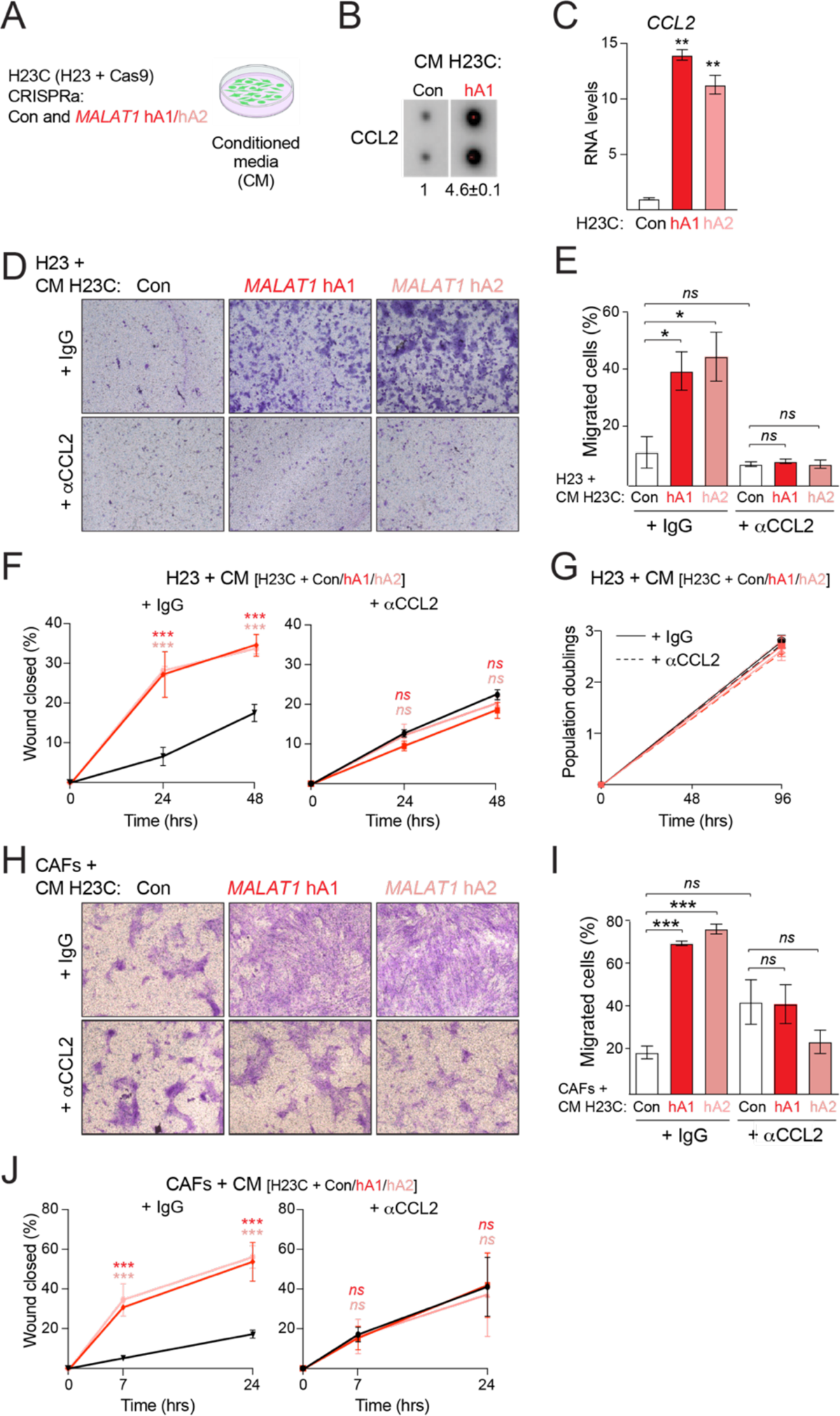
Ccl2 secretion mediates cellular phenotypes of *Malat1* overexpression. (A) CRISPRa of human *Malat1* in H23 cells, transduced with Cas9 (H23C); (B) Ccl2 protein levels by proteome profiling of CM from H23C+Con or H23C+hA1 cells (Fig. S5); (C) Ccl2 RNA levels in indicated H23C cells; (D) Representative images of transwell migration assay of H23C cells incubated with CM from indicated H23C cells in the absence (IgG) or presence of Ccl2-neutralizing antibody (αCcl2); (E) Quantification of transwell migration assay described in (D); (F) Quantification cell migration by wound healing assay of cells in (D); (G) Growth analysis of cells in (D); (H) Representative images of transwell migration assay of CAFs incubated with CM from indicated H23C cells in the absence (IgG) or presence of Ccl2-neutralizing antibody (α-Ccl2); (I) Quantification of transwell migration assay described in (H); (J) Wound healing assay of cells in (H). Data show mean ± SEM (n = 3, biological replicates).

To test the functional contribution of Ccl2 downstream of *Malat1* overexpression in LUAD, we examined how the presence or absence of a Ccl2-specific neutralizing antibody (αCcl2) affected cellular phenotypes of *Malat1* overexpression. Strikingly, we observed that addition of αCcl2 to the CM of *Malat1*-overexpressing H23C cells (H23C+hA1 and H23C+hA2) was sufficient to completely abrogate the *Malat1-*dependent increase in transwell migration (Fig. 6D, E) and wound healing (Fig. 6F) of tumor cells. Neutralization of Ccl2 did not significantly affect the behavior of H23C+Con cells, which lack *Malat1* overexpression (Fig. 6E, F), and did not impact cellular proliferation (Fig. 6G). Similarly, the addition of the Ccl2-neutralizing antibody to CM collected from *Malat1*-overexpressing H23C cells fully reversed the increased mobility of CAFs in response to *Malat1* overexpression while not impacting CAFs exposed to CM from H23C+Con cells (Fig. 6H, I, J). These epistasis experiments pointed to Ccl2 as the primary effector of cellular mobility downstream of overexpressed *Malat1*.

We also confirmed dramatic increases in murine Ccl2 protein and *Ccl2* transcript levels in CM and RNA, respectively, isolated from mA1 compared to a control KPC cell line (Fig. S6A, B, C). Analogous to human cells, the addition of the Ccl2 neutralizing antibody to CM from KPC mA1 cells also abrogated the *Malat1*-dependent increase in migration of tumor cells and CAFs, while addition of αCcl2 to CM from KPC Con cells did not alter cellular behavior (Fig. S6D, E). Inhibition of Ccl2 also did not affect the proliferation of murine KPC Con and mA1 cells (Fig. S6F). These results led us to conclude that secretion of Ccl2 is responsible for the increased cellular mobility downstream of *Malat1* overexpression.

### Ccl2 blockage suppresses inflammatory responses and tumor progression in *Malat1*-overexpressing LUAD

Ccl2 is an established mediator of breast and prostate cancer metastasis through its ability to recruit innate immune cells to the tumor microenvironment, including myeloid-derived suppressor cells and pro-tumor macrophages (Qian et al., 2011; Su et al., 2019). Moreover, Ccl2 inhibition via neutralizing antibodies has previously been pursued as a therapeutic strategy in metastatic breast and prostate cancer (Kirk et al., 2013; Qian *et al*., 2011).

To examine the contribution of Ccl2 to LUAD progression in *Malat1*-overexpressing KPC mice, we analyzed RNA sequencing data from Fig. 3A. We observed that while mature *Ccl2* was not detectable in control samples, it was expressed to high levels in all three *Malat1*-overexpressing KPC tumors (Fig. 7A). Consistently, we observed that *Malat1* overexpression correlated with a 5-fold increase in the recruitment of tumor-infiltrating macrophages in mA1 and mA2 KPC tumors compared to Con KPC tumors (Fig. 7B, C).

**Figure 7.**
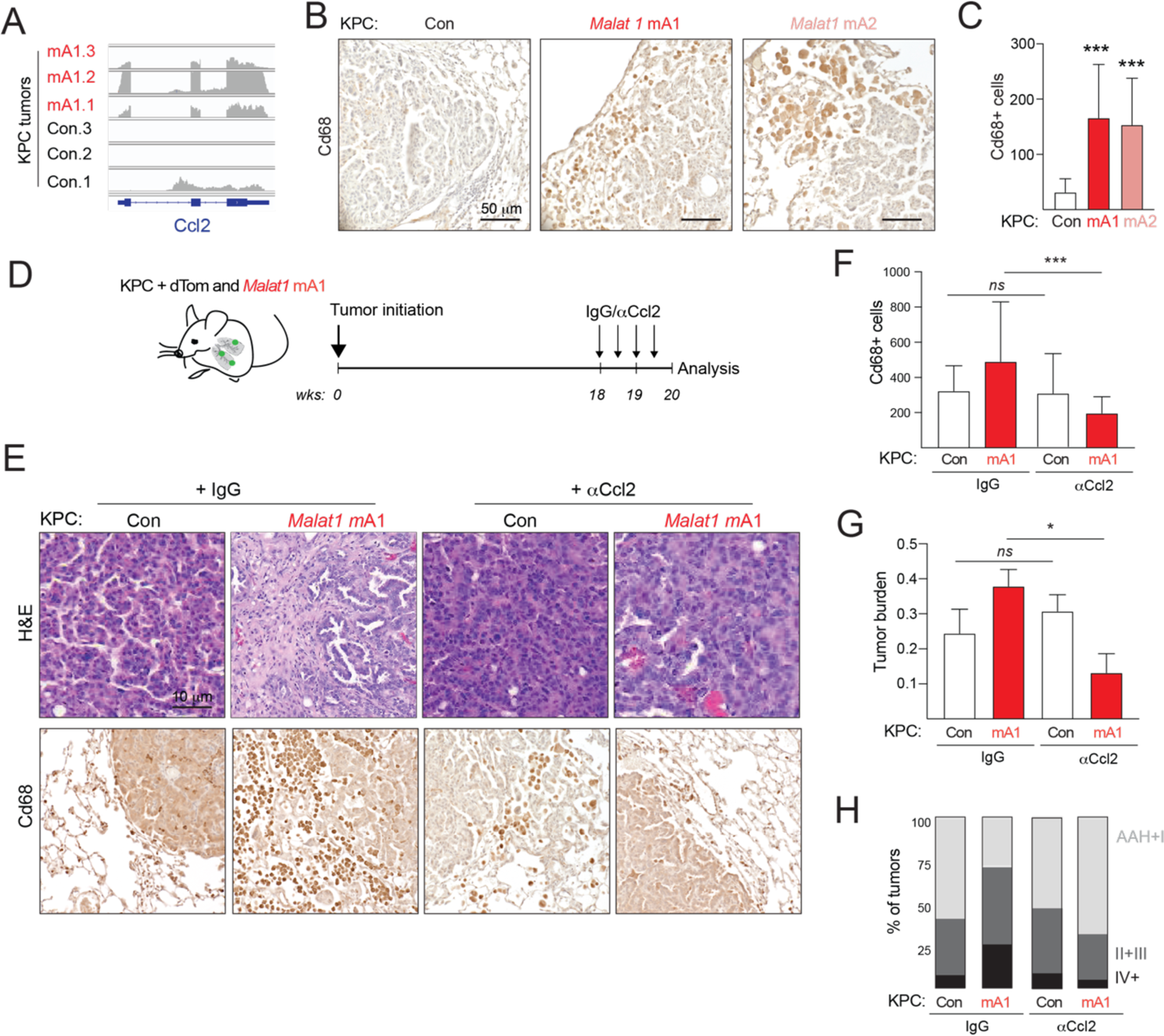
Ccl2 secretion establishes inflammatory tumor microenvironment and mediates tumor progression in *Malat1* overexpressing LUAD. (A) Read coverage of Ccl2 gene in RNA sequencing data in Con and mA1 KPC tumor cells; (B) Representative Cd68 IHC staining in tumor-bearing lungs from KPC mice infected with indicated dRNAs; (C) Quantification of Cd68 staining in (B); (D) Schema of *Malat1* CRISPRa in KPC animals followed by treatments with α-Ccl2 or IgG Con antibodies. Animals were treated biweekly at 18 and 19 weeks pti (n≥5 mice per experimental condition); (E) H&E and Cd68 IHC staining of lung sections of KPC mice infected and treated with indicated dRNAs and antibodies; (F) Quantification of Cd68 staining in (E); n=10 tumors per animal, unpaired t-test; (G) Quantification of tumor burden in mice in (D); (H) Quantification of tumor grade in mice in (D).

These data led us to hypothesize that inappropriate secretion of Ccl2 by tumor cells may be directly responsible for increased macrophage recruitment in *Malat1* overexpressing tumors and that blocking Ccl2 activity in the context of *Malat1* overexpression may reverse phenotypes of tumor progression *in vivo*. To test these hypotheses, we assembled a cohort of Con- and mA1-expressing KPC animals at 18 weeks post tumor initiation and administered four biweekly intraperitoneal injections with control IgG or αCcl2 antibodies (Fig. 7D). Immunohistochemistry staining at experimental endpoint revealed that inhibition of Ccl2 did not have a significant effect on the recruitment of tumor-infiltrating macrophages in Con tumors but suppressed the excessive macrophage infiltration in *Malat1*-overexpressing tumors (Fig. 7E, F). In addition to changes in the tumor microenvironment, Ccl2 blockade led to a significant reduction of both tumor burden and grade in *Malat1*-overexpressing but not control KPC mice (Fig. 7G, H). We concluded that secreted Ccl2 is the primary paracrine effector of macrophage recruitment downstream of *Malat1* overexpression *in vivo* and that inhibition of Ccl2 activity reverses key phenotypes of tumor progression in *Malat1*-overexpressing advanced LUAD.

## Discussion

While protein-coding determinants of metastatic cancer have been extensively studied, lncRNAs that contribute to metastatic progression are less well characterized and frequently lack experimental validation in relevant cancer models (Olivero and Dimitrova, 2020). In this study, we present conclusive evidence that the lncRNA *Malat1* is a *bona fide* proto-oncogene and that *Malat1* overexpression is sufficient to overcome the barrier to metastatic progression in the KP mouse model of LUAD.

The robust correlation between *Malat1* overexpression and metastatic progression in LUAD patients was initially observed almost two decades ago (Ji *et al*., 2003; Muller-Tidow *et al*., 2004). Yet efforts to model *Malat1* overexpression in immunocompetent LUAD models have lagged and the role of *Malat1* in shaping LUAD evolution has remained poorly understood (Arun *et al*., 2020; Sun and Ma, 2019). In this study, we describe the first successful approach to carry out *Malat1* overexpression in the KP LUAD mouse model, which closely recapitulates the progression of human LUAD including the sporadic evolution to metastatic disease. Notably, CRISPRa of *Malat1* in KP lung tumors led to tumor-specific overexpression of endogenous *Malat1* and resulted in the synchronous development of control and *Malat1*-overexpressing tumors in an immunocompetent host. Our results revealed that *Malat1* overexpression is a powerful driver of tumor progression to a poorly differentiated, aggressive, and metastatic disease. Therefore, *Malat1* is not simply a prognostic marker but an oncogenic driver of metastatic progression. Interestingly, we observed that *Malat1* overexpression only had phenotypic consequences in the context of p53 deficiency, indicating that *Malat1* overexpression acts in cooperation with p53 loss to promote tumor progression and that p53 function antagonizes the consequences of *Malat1* overexpression. In sum, modeling of *Malat1* overexpression *in vivo* showed a driver role for this lncRNA in LUAD progression and provided novel insights into the interplay between genetic and epigenetic determinants of metastatic dissemination in LUAD.

These findings are consistent with a body of work that has ascribed oncogenic functions to *Malat1* but in contrast to studies that have reported tumor suppressive activities (Arun *et al*., 2020; Sun and Ma, 2019). We propose that context-specific differences may explain the discrepancies, such as the status of the p53 pathway or the specifics of the *Malat1* perturbation model, which may not accurately recapitulate the cancer-associated overexpression of *Malat1*.

Interestingly, we observed a widespread increase of *Malat1* levels over time in the synchronously progressing KP tumors, analogous to the overexpression of *Malat1* in advanced human LUAD. This points to a conserved mechanism that links tumor progression with intrinsic *Malat1* overexpression in human and mouse cancer. Moreover, we identified the inflammatory cytokine Ccl2 as the principal paracrine effector downstream of overexpressed *Malat1* in both human and murine LUAD models. Thus, the oncogenic roles of *Malat1* overexpression are conserved between mouse and human, and insights from the murine model are likely to be relevant to human cancer.

Our work indicates that *Malat1* overexpression in cancer is a double-edged sword that cooperatively activates tumor cell autonomous and non-cell autonomous mechanisms to promote metastatic reprogramming. On the one hand, we report that *Malat1* overexpression alters the differentiation state of tumor cells. The observed loss of epithelial identify, EMT, and increased cellular mobility of tumor cells are established hallmarks of metastatic progression that have previously been associated with *Malat1* overexpression (Arun *et al*., 2016; Guo *et al*., 2015; Gutschner *et al*., 2013; Tano *et al*., 2010). On the other hand, we uncover an unexpected link between *Malat1* overexpression and activation of innate immune signaling through inappropriate expression and secretion of the inflammatory cytokine Ccl2. The innate immune system represents the first line of defense to foreign substances. However, innate immune cells, such as pro-tumor macrophages, recruited by Ccl2 signaling have also been shown to drive cancer-associated inflammation, and to promote steps of the metastatic cascade, including ECM remodeling, vascularization, and cancer cell mobility (Lim et al., 2016). Indeed, we observed that *Malat1* overexpression is associated with Ccl2-dependent recruitment of macrophages, pleiotropic changes in the tumor stroma, and enhanced cancer and stromal cell mobility. We presented evidence that addition of a Ccl2-neutralizing antibody to conditioned media from *Malat1*-overexpressing human and mouse LUAD cells abolished the enhanced mobility of tumor and stromal cells. Moreover, treatment of tumor-bearing *Malat1*-overexpressing KP tumors with the same Ccl2-neutralizing antibody reversed macrophage recruitment and rescued stromal remodeling. Together, these experiments establish that overexpressed *Malat1* acts through Ccl2 signaling to promote cancer progression and that the consequences of *Malat1* overexpression can be reversed by inhibition of Ccl2 signaling.

We propose that the consequences of overexpressed *Malat1* represent novel gain-of-function activities rather than perturbations of its physiological functions. There is evidence that loss of *Malat1* in a breast carcinoma model prevents dedifferentiation of mammary tumors (Arun *et al*., 2016). Similarly, we observed that *Malat1* overexpression by CRISPRa correlated with LUAD dedifferentiation. Thus, it appears that overexpression of *Malat1* resulting from or accompanying tumor progression promotes loss of epithelial identify. However, epithelial differentiation was not found to be perturbed in *Malat1*-deficient mice (Eissmann *et al*., 2012; Nakagawa *et al*., 2012; Zhang *et al*., 2012), suggesting that epithelial dedifferentiation is an overexpression-specific phenotype. Analogously, Ccl2 is not normally expressed and secreted by epithelial cells but becomes inappropriately turned on in tumor cells in the context of *Malat1* overexpression. In both cases, we determined that gene expression changes following *Malat1* overexpression were due to transcriptional deregulation rather than as a result of aberrations in alternative splicing or post-transcriptional control.

The observed centrality of Ccl2 in promoting immune evasion and tumor progression in KP LUAD is consistent with previous studies (Martin et al., 2021; Su *et al*., 2019). Our study further adds *Malat1* overexpression as a novel and potentially widespread route for Ccl2 activation in cancer. Given the recurrent overexpression of *Malat1* across many cancer types, it is tempting to speculate that *Malat1* overexpression in pre-metastatic disease may serve as a valuable prognostic marker for Ccl2 therapy. Conversely, given the risks of Ccl2 therapy (Bonapace et al., 2014), targeting *Malat1* via antisense oligonucleotides (ASOs) (Arun *et al*., 2016; Gutschner *et al*., 2013) or ENE-targeting compounds (Abulwerdi et al., 2019; Zafferani et al., 2022) may be an attractive alternative due lack of anticipated side-effects. Finally, it will also be important to determine whether *Malat1* overexpression is a marker of immunosuppression and whether *Malat1* inhibition may improve the performance of immune checkpoint inhibitors.

## Materials and Methods

### Mouse strains

All animal work was conducted in accordance with a protocol approved by the Yale University Institutional Animal Care and Use Committee. K-rasLSL-G12D/+ (K) and p53flox/flox (P) mice were previously described and obtained from the laboratory of T. Jacks (MIT) (Jackson *et al*., 2005; Jackson *et al*., 2001). Rosa26-Cas9-T2A-GFPLSL/LSL (C) (strain #027619, (Platt *et al*., 2014)) was purchased from Jackson Laboratories. For tumor studies, 3-6 months-old mice were used. Experiments were performed blind to gender and with an equal distribution of males and females in each experimental group.

### Cell culture

Human lung adenocarcinoma cell lines H23 (American Type Culture Collection (ATCC)), H2009 (ATCC) and H2122 (German Collection of Microorganisms and Cell Culture (DMSZ)) were grown in RPMI 1640 (Thermo Scientific) supplemented with 10% FBS (F0926, Sigma-Aldrich), 50 U/ml pen/strep (GIBCO), and 2 mM L-glutamine (GIBCO). CAFs were purchased from Neuromics and grown in VitroPlus III, low serum, complete medium (Neuromics). KPC cells were derived from tumor-bearing KPC lungs dissociated with collagenase for 30 min at 37°C. Murine tumor cell lines were maintained in DMEM (GIBCO) supplemented with 10-15% FBS (F0926, Sigma-Aldrich), 50 U/ml pen/strep (GIBCO), 2 mM L-glutamine (GIBCO), 0.1 mM non-essential amino acids (GIBCO), and 0.055 mM β-mercaptoethanol (GIBCO). Cell lines tested negative for mycoplasma using MycoAlert (Lonza: LT07-28). All experiments were performed within 10 passages after thawing cells. Genotypes were confirmed by genotyping and by qRT-PCR. 293 viral packaging cells were cultured in DMEM supplemented with 10% FBS, 50 U/ml pen/strep, 2 mM L-glutamine, and 0.1 mM non-essential amino acids. All cell cultures were maintained at 37C in a humidified incubator with 5% CO2.

### Constructs

In vitro CRISPRa experiments were performed with a previously described lentiviral vector (lenti-SAM-Hygro (Olivero et al., 2020)). In vitro CRISPRi experiments were performed with a lentiviral vector (lenti-KRAB-Hygro) constructed to co-express a U6-driven 15-mer ʻdead RNA’ (dRNA) extended by two MS2 loops (dRNA-MS2), the transcriptional repressor KRAB (Kruppel-associated Box) fused to the MS2-binding protein (MBP), and a hygromycin-resistance gene. CRISPRa and CRISPRi constructs were introduced in cell lines previously transduced with spCas9 from SpCas9-T2A-Puro (BRD001, gift from the Broad Institute, MIT) construct introduced in KP cells and selected with puromycin treatment (5 µg/ml, Sigma-Aldrich)) or SpCas9-T2A-GFP (BRD004, gift from the Broad Institute, MIT) construct introduced in H23, H2009, and H2122 cells and selected by GFP+ FACS sorting). CRISPRa- and CRISPRi-expressing cells were selected with hygromycin (0.8 µg/ml, Roche). Lentivirus was produced in 293 cells by co-transfecting the lentiviral constructs with pCMV-dR8.2 (Addgene plasmid #8455) and pCMV-VSV-G (Addgene plasmid #8454) viral packaging constructs. Viral containing supernatants supplemented with 4 µg/ml polybrene (Millipore Sigma) were used to infect the cells by 2 consecutive lentiviral infections, delivered at 24 hour-intervals.

For CRISPRa *in vivo*, dRNAs were cloned downstream of a U6 promoter in Lenti-dRNA-SAM-Cre lentiviral vector, engineered analogously as the in vitro system. Lentivirus was prepared as above, concentrated by ultracentrifugation, and titered by infecting 3TZ cells, a derivative of 3T3 cells, expressing a LSL-LacZ transgene (generously provided by the laboratory of T. Jacks, MIT). Titer was determined as the number of viral particles based on the fraction of LacZ-positive cells as previously described (DuPage *et al*., 2009). Mutagenesis of the ENE domain in cultured cells was performed with a sgRNA targeting the ENE domain of *Malat1*, expressed from the BRD001 lentiviral construct. To determine mutagenesis efficiency, a region spanning the ENE domain was amplified by conventional PCR (PrimeStar HS mix, Takara Bio) with primer sequences listed in Table S9. The amplicon was cloned into pCR-Blunt II-TOPO® vector (Invitrogen) and analyzed by Sanger sequencing. All dRNA and sgRNA sequences are listed in Table S9.

### Patient samples

Formalin-fixed paraffin-embedded (FFPE) samples were collected, processed, and represented in tissue microarray (TMA) by Yale Pathology Tissue Services (YPTS). The cohort (YTMA) was composed of 161 NSCLC de-identified lung cancer patients enrolled between 2011-2016. All cases in the cohort were reviewed by a local pathologist using H&E-stained preparations and tumor histology variant was confirmed by morphology analysis. Tumor cores for TMA construction were obtained from case areas selected by a pathologist to represent the disease. Patient clinicopathologic information was collected from clinical records and pathology reports. Tissues were collected in accordance with consent guidelines from the Yale Human Investigation Committee.

### RNA isolation and RT-qPCR

For RNA-seq and RT-qPCR analysis, RNA was treated with DNaseI (Qiagen) and isolated with the RNeasy Mini Kit (Qiagen). 0.5-1 μg of total RNA was reverse transcribed using the High-Capacity cDNA Reverse Transcription Kit (Applied Biosystems). SYBR Green PCR master mix (Kapa Biosystems) was used for quantitative PCR in triplicate reactions with primers listed in Table S9. RNA expression levels were calculated using the ddCt method relative to GAPDH and normalized to control samples.

### Antisense oligonucleotides (ASOs)

1 × 10^5^ *Malat1*-overexpressing H23+hA1 and H23+hA2 cells, seeded in 6-well plates, were transfected with 10 nM *Malat1* ENE-targeting (ASO) or control (Con) steric blocking ASOs (Exiqon, Qiagen) using Lipofectamine 2000 (ThermoFisher Scientific). *Malat1* RNA levels and the corresponding effects on cellular migration were assayed at 72 hours post-transfection by qRT-PCR and Boyden Chamber assay. The sequences of the ASOs are listed in Table S9.

### Cellular assays

Cellular proliferation was assessed by plating 1 × 10^5^ cells in 6-cm plates. Cells were harvested and counted at indicated times, and cumulative cell numbers were plotted over time.

Wound healing assay was performed by scraping a confluent monolayer of cells grown in 6-well plates with a P-200 micropipette tip. Media was then changed by adding new complete RPMI media or CM collected and filtered (0.2 µm) from indicated cells. Pictures of the wounds were taken at the indicated times. The area of the wound healing was measured with Fiji software. Three wells and three measures/well were used per condition and experiment. Transwell migration, also known as Boyden Chamber, assay was performed with 1 × 10^5^ cells resuspended in serum-free RPMI media added to the upper chamber of 8.0-µm-pore transwell filters (Corning, 3422). For conditioned media (CM) experiments, cells were pre-incubated with CM from indicated cells for 48 hours prior to seeding. The lower chamber contained complete RPMI media or CM collected from indicated cells and passed through a 0.2 µm filter, supplemented with 1 µg/ml CCL2 antibody (BioCell, BE0185) or isotype control IgG (BioCell, BE0091), as indicated. After 24 hours incubation at 37°C, inserts were fixed in formalin 4%, and stained in 0.2% crystal violet. Pictures were analyzed using Fiji software.

### Tumor studies

Lung tumorigenesis was initiated in cohorts of KC/KPC mice as described in (DuPage *et al*., 2009) by intratracheal infection with 1 x 10^5^ lentiviral particles. Mice were analyzed at 16 weeks post tumor initiation (pti). For histological analyses, tumor-bearing lungs were inflated with 4% paraformaldehyde, and fixed in 4% paraformaldehyde for 24 hours, prior to dehydration in 70% ethanol. Fixed lungs were embedded in paraffin, sectioned, and stained with hematoxylin and eosin (H&E) or Masson’s trichrome. Images were acquired with an Axio Imager 2 microscope system (Zeiss) with a PlanApo 10x 0.3 objective lens (Zeiss). Tumor burden, scored as the fraction of total lung area, was determined using the freehand selection tool and Measure feature in ImageJ, and normalized to control animals. Tumor grade was scored as previously described (DuPage *et al*., 2009; Nikitin et al., 2004). In short, Advanced Adenomatous Hyperplasia (AAH): peripheral focal proliferation of atypical cuboidal epithelial cells, adjacent to normal lung; grade 1: tumors with uniform nuclei showing no nuclear atypia; grade 2: tumors containing cells with uniform but slightly enlarged nuclei that exhibit prominent nucleoli; grade 3: tumor cells with enlarged, pleomorphic nuclei showing prominent nucleoli and nuclear molding; grade 4: tumor cells showing large, pleomorphic nuclei exhibiting a high degree of nuclear atypia, including abnormal mitoses and hyperchromatism, and containing multinucleate giant cells; grade 5: tumors with all the features of grade 4 tumors and also showing stromal desmoplasia surrounding nests of tumor cells.

Overall survival was evaluated by comparing the survival of *Malat1*-overexpressing and control mice for a duration of 25 weeks from tumor initiation. Animals were euthanized at ethical endpoint criteria, including significant body weight loss, hunched posture, or impaired mobility. Macroscopic and microscopic primary and metastatic tumors were confirmed by GFP signal under an inverted microscope. Twice a week treatments with the CCL2 neutralizing antibody or IgG control were performed in randomized mA1- or control-expressing groups starting at 18 weeks pti. Each mouse was given a 100-μl intraperitoneal injection of 200 μg of each antibody diluted in PBS.

### Immunohistochemistry

Indirect immunohistochemistry was carried out on paraffin-embedded tissue sections using the signal amplification ABC Vectastain kit (Vector Labs). Endogenous peroxidase activity was blocked with 3% hydrogen peroxidase for 10 min and antigen retrieval was carried out by heating sections in a steamer with citrate buffer (pH 6) for 30 min at 95°C. Afterwards, tissues were incubated with primary antibodies: Hmga2 (BioChek, 59170AP; 1:500), GFP (Cell Signaling Technology, 2555S; 1:2000), α-Sma (ThermoScientific, 14-9760-82; 1:000), Cd68 (Abcam, ab125212; 1:100), Ki67 (D3B5, Rabbit mAb Mouse Preferred IHC Formulated; 1:500), pHH3 (Cell Signaling Technology, 9701S; 1:500), and Cdh1 (Cell Signaling, 24E10, 1:1000) at 4°C overnight. The signal was visualized with DAB (Vector Labs) and slides were counterstained with hematoxylin. Hmga2 and GFP staining was evaluated by qualitative interpretation as negative (≤5% positive cells) or positive (>5% positive cells) staining. Ki67, pHH3, and Cd68 staining was evaluated as the percentage of positive cells per tumor using automated QuPath software (Bankhead et al., 2017). α-Sma and Cdh1 staining was assessed by H-Score semi-quantitative method as previously described (Martinez-Terroba et al., 2018).

### RNAScope

Detection of *Malat1* expression levels (Love et al., 2014) was examined in lung cancer patient cohort TMA and murine lung tumor sections, respectively, using an RNAScope 2.5 HD Reagent Kit-BROWN (322300) and probes (human *Malat1*, ACD Bio-Techne, #400811; mouse *Malat1*, ACD Bio-Techne, #313391) following manufacturer instructions. Briefly, slides were baked at 60°C for 30 min using the HybEZ Oven, prior to pretreating with pretreat 2 and Protease Plus for 5 min at 95°C and 40°C, respectively. Following amplification steps, sections were incubated for 10 seconds in DAB solution and then washed with water and counterstained. Signal was evaluated by Qupath software (Quantitative Pathology & Bioimage Analysis) (Bankhead *et al*., 2017) as well as H-Score method as previously described (Martinez-Terroba *et al*., 2018).

### Flow cytometry and cell sorting

Lung tumors were minced using a razor blade and incubated at 37°C for 30 min in dissociation solution containing DNAse (1µg/ml, Sigma-Aldrich) and Collagenase IV (200 U/ml, Sigma-Aldrich). Following red blood cell lysis (ACK lysis buffer, Lonza), single cell suspensions of lung tumors were resuspended in FACS buffer (PBS, 1% FBS). Cells were incubated with anti-Fc receptor antibody (Biolegend, 101320) on ice for 15 min followed by staining with the following antibodies for 30 min: 1:500 CD45, Pacific Blue (BioLegend, 10325); 1:500 Live/Dead Red Dye (Invitrogen); 1:500 CD326-EpCAM, APC (Biolegend, 118211). Live/ GFP+ /CD45-/ Epcam+ tumor epithelial cells were sorted on a BD FACS Aria at the Yale Flow Cytometry Core facility.

### RNA sequencing

Total RNA was isolated from sorted tumor cells (Live/GFP+/CD45-/ Epcam+) in three biological replicates from different animals infected with Con or mA1 dRNAs at 16 weeks post tumor initiation. RNA was treated with DNaseI (Qiagen) and isolated with the RNeasy Micro Kit (Qiagen). PolyA selection, cDNA library preparation using NEBNext Single Cell/Low Input RNA Library Prep Kit for Illumina (NEB), and sequencing was performed at the Yale Stem Cell Center Genomics Core facility. mRNA-seq reads were mapped to the mm10 genome assembly (Schneider et al., 2017) with TopHat (2.1.1) (Trapnell et al., 2009). GENCODE transcript annotation (M9) (Frankish et al., 2019) was used to determine nonoverlapping exons or intronic regions using BEDTools (Quinlan and Hall, 2010) and custom scripts. The generated exonic or intronic annotations were used to determine overlapping sequencing reads that were counted with HTSeq (1.99.2) (Anders et al., 2015) and summed to determine gene levels with custom scripts. Annotated genes from chrX and chrY were excluded from the analysis. PCA was done in R and plots generated with ggplot2. Gene sets from the Molecular Signature Database (mSigDB) (Liberzon et al., 2015) were downloaded from the GSEA webpage (http://software.broadinstitute.org/gsea) (Subramanian et al., 2005). Differentially expressed genes were determined using DESeq2 (Love *et al*., 2014) with an expression cutoff of >10 normalized reads in any two of the six samples.

### Antibody arrays

Cytokines, chemokines, and growth factors secreted by indicated cells were detected in conditioned media (CM) using RayBio Human Angiogenesis Antibody array C1000 (RayBiotech) and Proteome Profiler Human Protease Array Kit (R&D systems) according to the manufacturer’s instructions. The signal intensity of each spot was evaluated and normalized to positive controls using ImageJ software.

### ELISA

Ccl2 protein levels were measured in 0.2 µm-filtered CM, collected from indicated cells, with the Mouse Monocyte Chemoattractant Protein-1/CCL2 (MCP-1) Uncoated ELISA Kit (ThermoFisher) according to manufacturer’s instructions. Briefly, 96-well plates were coated with Ccl2 antibody (1:250) prior to blocking with ELISA/ELISPOT diluent. Samples and standards, consisting of serial dilutions of murine Ccl2 reconstituted in distilled water, were added to the plates and detection was performed using a biotin-conjugated anti-mouse Ccl2 antibody, diluted 1:250 in ELISA/ELISPOT diluent, followed by incubation with Streptavidin-HRP, diluted 1:100 in ELISA/ELISPOT diluent. Tetramethylbenzidine substrate solution was added to the enzyme-antibody-target complexes to produce measurable signal and reactions were stopped by adding 2N H2SO4. Measurements at 450 nm relative to 570 nm were performed on a SpectraMax iD3 microplate reader.

### Statistical analysis

Student’s t-test (paired and unpaired, as appropriate) and non-parametric Mann-Whitney U test (two-tailed, unpaired) were used for statistical analysis of the results of cellular proliferation, cellular migration, RNA levels, and tumor burden after testing for normality. Wilcoxon signed rank sum test was applied to analyze *Malat1* differences between tumor and normal adjacent tissue. Pearson’s Chi-square test and Fisher’s exact test were used to analyze differences in *Malat1* RNA levels between two or more than two groups, following Cochran recommendations. For survival analysis, patients were stratified into two groups as indicated according to *Malat1* expression levels. Cumulative survival of patients was estimated using Kaplan-Meier curves, and significant differences between groups were tested using the log-rank test. The influence of clinicopathologic variables on recurrence was assessed with the Cox proportional hazards model. Disease-free survival was calculated from the date of surgery to the date of recurrence or last follow-up. The follow-up period was restricted to 60 months. All statistical analyses were performed, and graphics were generated using Prism8 software. ns, not significant; ∗p <0.05, ∗∗p < 0.01, and ∗∗∗p < 0.001.

## Acknowledgements

We thank Dr. David Rimm from the Yale Pathology Tissue Service, the Yale Center for Genome Analysis, the Keck DNA Sequencing Facility, and the Yale Flow Cytometry Facility for materials and analysis support. This work was supported in part by the LCRF (ND), the V Foundation (ND), the Pew-Stewart Scholars Program for Cancer Research (ND), NIH R37CA230580 (ND), NIH RO1CA262286 (ND), NIH R01GM143536 (JRZ), by Developmental Research Project Awards funded by Yale SPORE in Lung Cancer NIH P50CA196530 (ND, KP) and by Yale Cancer Center Support Grant NIH P30CA016359 (ND, KP). EMT was supported by an American Cancer Society Postdoctoral Fellowship (PF-20-103-01).

## Author contributions

EMT and ND conceived the project, analyzed data, prepared figures, and wrote the manuscript. EMT performed experiments. FJM and CRO provided experimental assistance with *in vivo* studies and VL and JRZ performed transcriptome analysis. ND, JRZ, and KP supervised the project.

## Competing Interest Statement

The authors declare no competing interests

## Notes

### Competing Interest Statement

The authors have declared no competing interest.

