## Supplementary Figures and Legends for "Overexpressed *Malat1* Drives Metastasis through Inflammatory Reprogramming of Lung Adenocarcinoma Microenvironment"

### Supplementary Information

#### Supplementary text

#### Supplementary tables S1-S9

#### Supplementary figures and figure legends S1-S6

**Supplementary text**

To determine the clinical significance of *Malat1* expression in lung cancer, we evaluated the expression of *Malat1* by RNAScope technology in formalin-fixed paraffin embedded (FFPE) tissue microarray (TMA) of samples from a cohort of lung cancer tumors (n=161). Probe specificity and reproducibility validation is shown in Fig. S1B-C. Clinical data from these patients are listed in Table S1. Normal adjacent tissue, including pneumocytes, macrophages, stromal and epithelial cells from bronchioles from the alveolar parenchyma exhibit ubiquitous *Malat1* expression in the nucleus, as expected. However, *Malat1* expression was significantly elevated in tumor tissue compared to normal adjacent tissue (Figs. 1A and B; $p<0.001$). Evaluation of the relationship between *Malat1* levels and clinicopathological features revealed lack of correlation between *Malat1* expression and age, gender, smoking status or stage (Table S2). A significant correlation was found between *Malat1* RNA levels and histology. Specifically, *Malat1* levels were significantly elevated in lung adenocarcinoma (LUAD) compared to squamous cell carcinoma (SCC) (Table S2). Next, we investigated whether the expression of *Malat1* was associated with prognosis. Patients were stratified in two groups (lower tertile and top two tertiles) according to *Malat1* expression and differences were evaluated using log-rank test. High *Malat1* RNA levels were significantly associated with a decrease in disease-free time (Fig. 1C; $p=0.012$). A multivariate Cox regression analysis, adjusted by stage, showed that *Malat1* expression is an independent prognostic factor for disease-free survival ($p=0.021$, HR = 1.132; 95% CI, 1.187-8.109, Table S3). Altogether, these results support the conclusion that high expression of *Malat1* is associated with poor prognosis in lung cancer.

**Supplementary tables**

**Table S1.** Clinicopathological characteristics of the NSCLC patients included in the study

| Characteristics | YTMA cohort (n=161) |
|---|---|
| Age | |
| ≤65 | 69 (42.9%) |
| >65 | 92 (57.1%) |
| Gender | |
| Male | 61 (37.9%) |
| Female | 100 (62.1%) |
| Smoking status | |
| Never | 18 (11.2%) |
| Former | 103 (64.0%) |
| Current | 40 (24.8%) |
| Histology | |
| LUAD | 108 (67.0%) |
| SCC | 45 (28.0%) |
| Others | 8 (5.0%) |
| Stage | |
| I | 132 (82.0%) |
| II-IV | 29 (18.0%) |

**Table S2.** Relationship between clinicopathological characteristics and *MALAT1* expression in NSCLC patients

| Characteristics | YTMA cohort | | |
| --- | --- | --- | --- |
| | Low MALAT1 | High MALAT1 | P value |
| Age | | | |
| ≤65 | 27 (16.8%) | 42 (26.1%) | 0.316 |
| >65 | 29 (18.0%) | 63 (39.1%) | |
| Gender | | | |
| Male | 25 (15.5%) | 36 (22.4%) | 0.197 |
| Female | 31 (19.2%) | 69 (42.9%) | |
| Smoking status | | | |
| Never | 7 (4.3%) | 11 (6.8%) | 0.530 |
| Former | 38 (23.6%) | 65 (40.5%) | |
| Current | 11 (6.8%) | 29 (18.0%) | |
| Histology | | | |
| LUAD | 26 (16.1%) | 82 (50.9%) | **<0.001** |
| SCC | 28 (17.4%) | 17 (10.6%) | |
| Others | 2 (1.2%) | 6 (3.8%) | |
| Stage | | | |
| I | 47 (29.2%) | 85 (52.8%) | 0.640 |
| II-IV | 9 (5.6%) | 20 (12.4%) | |

**Table S3.** Univariate and multivariate Cox proportional hazards analysis for *MALAT1* expression and clinicopathological parameters in patients from the YTMA cohort

| | n (%) | Disease-free survival | | | |
|---|---|---|---|---|---|
| | | Univariate analysis | | Multivariate analysis | |
| | | HR (95% CI) | P value | HR (95% CI) | P value |
| MALAT1 | | | | | |
| Low | 56 (34.8%) | 1 | **0.018** | 1 | **0.021** |
| High | 105 (65.2%) | 1.161 (1.222-8.334) | | 1.132 (1.187-8.109) | |
| Age | | | | | |
| ≤65 | 69 (42.9%) | 1 | 0.538 | | |
| >65 | 92 (57.1%) | 0.227 (0.607-2.586) | | | |
| Gender | | | | | |
| Male | 61 (37.9%) | 1 | 0.309 | | |
| Female | 100 (62.1%) | 0.403 (0.689-3.251) | | | |
| Smoking status | | | | | |
| Never | 18 (11.2%) | 1 | 0.938 | | |
| Former | 103 (64.0%) | 0.011 (0.304-3.357) | | | |
| Current | 40 (24.8%) | 0.128 (0.385-2.009) | | | |
| Histology | | | | | |
| ADC | 108 (67.0%) | 1 | 0.511 | | |
| SCC | 45 (28.0%) | -0.214 (0.190-3.427) | | | |
| Others | 8 (5.0%) | -0.707 (0.099-2.446) | | | |
| Stage | | | | | |
| I | 132 (82.0%) | 1 | **0.156** | 1 | 0.212 |
| II-IV | 29 (18.0%) | 0.584 (0.88-4.018) | | 0.513 (0.746-3.745) | |

**Table S4.** Top 200 genes in PC1 in mA1/Con KPC samples

**Table S5.** DESeq2 normalized total read counts and summary from genes in Con and mA1 KPC samples

**Table S6.** Significantly upregulated and downregulated genes in mA1/Con KPC samples

**Table S7.** Core enrichment in Hallmark_EMT geneset upregulated with mA1

**Table S8.** DESeq2 normalized intronic read counts and summary from genes in mA1/Con

**Table S9.** Oligonucleotide, dRNA and sgRNA sequences used in this study.

**Supplementary Figures**

Martinez-Terroba et al., Figure S1

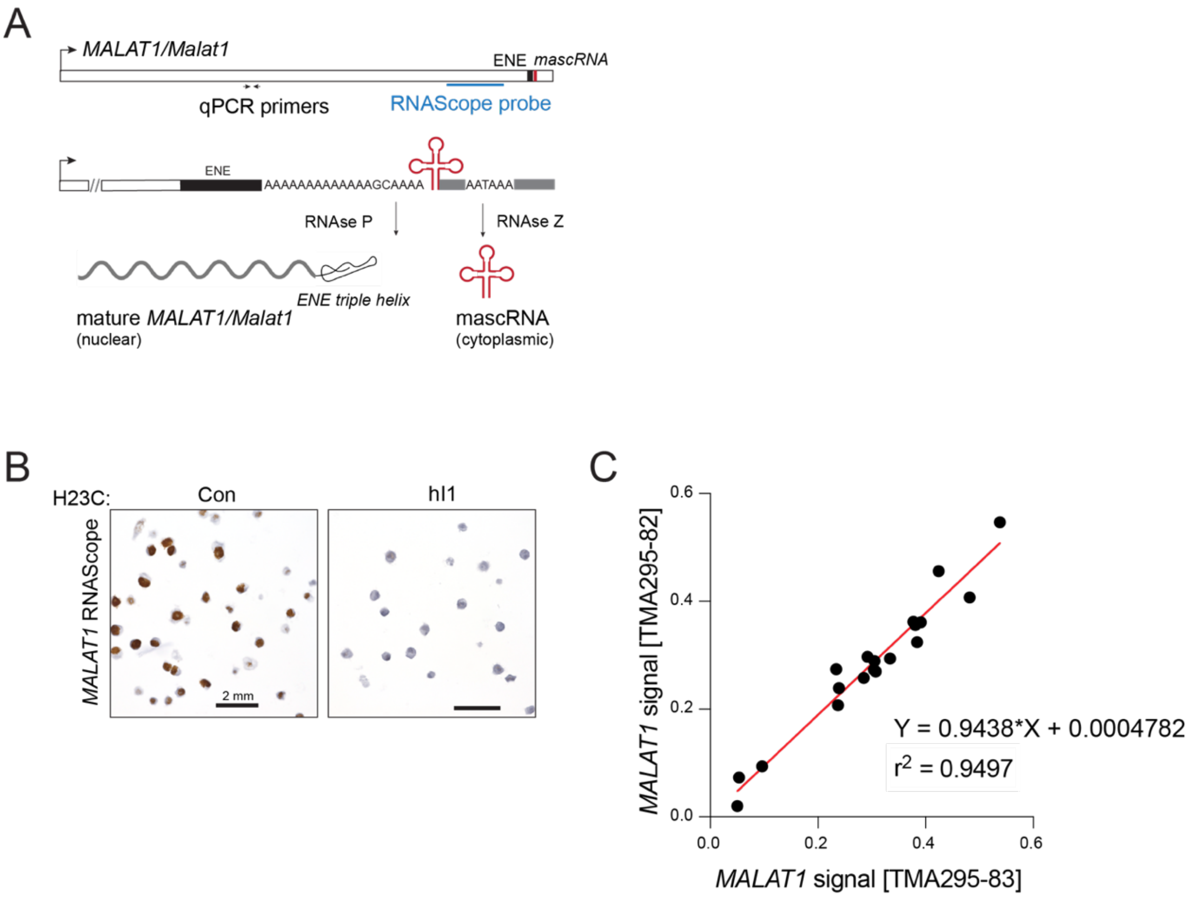


**<u>Figure S1.</u> Schematic of *Malat1* transcript processing and evaluation of the specificity and reproducibility of *Malat1* RNAScope probe.** (A) Schematic of *Malat1* unprocessed transcript highlighting ENE domain (black), tRNA-like mascRNA (red), and the locations of RT-qPCR primers (arrows) and RNAScope probe (blue line). The 3' end of *Malat1* is recognized and cleaved by RNase P. Despite lacking a poly(A) tail, *Malat1* RNA is protected from 3'-5' exonucleases by the formation of a conserved triple helical structure encoded by the ENE element; (B) Validation of *Malat1* probe specificity for RNAScope assay. RNAScope was performed in parafin-embedded H23C cells in the absence (Con) or presence (hI1) of *Malat1* CRISPRi. Transcriptional repression of *Malat1* cells led to loss of *Malat1* signal, verifying the probe specificity; (C) Validation of *Malat1* probe reproducibility. RNAScope was performed in two serial slides of TMAs on different days. *Malat1* nuclear signal was quantified using QuPath software. Correlation graph of *Malat1* signal in the serial TMA slides validate high *Malat1* probe performance (n=19 tumors, r2=0.9497).

Martinez-Terroba et al., Figure S2

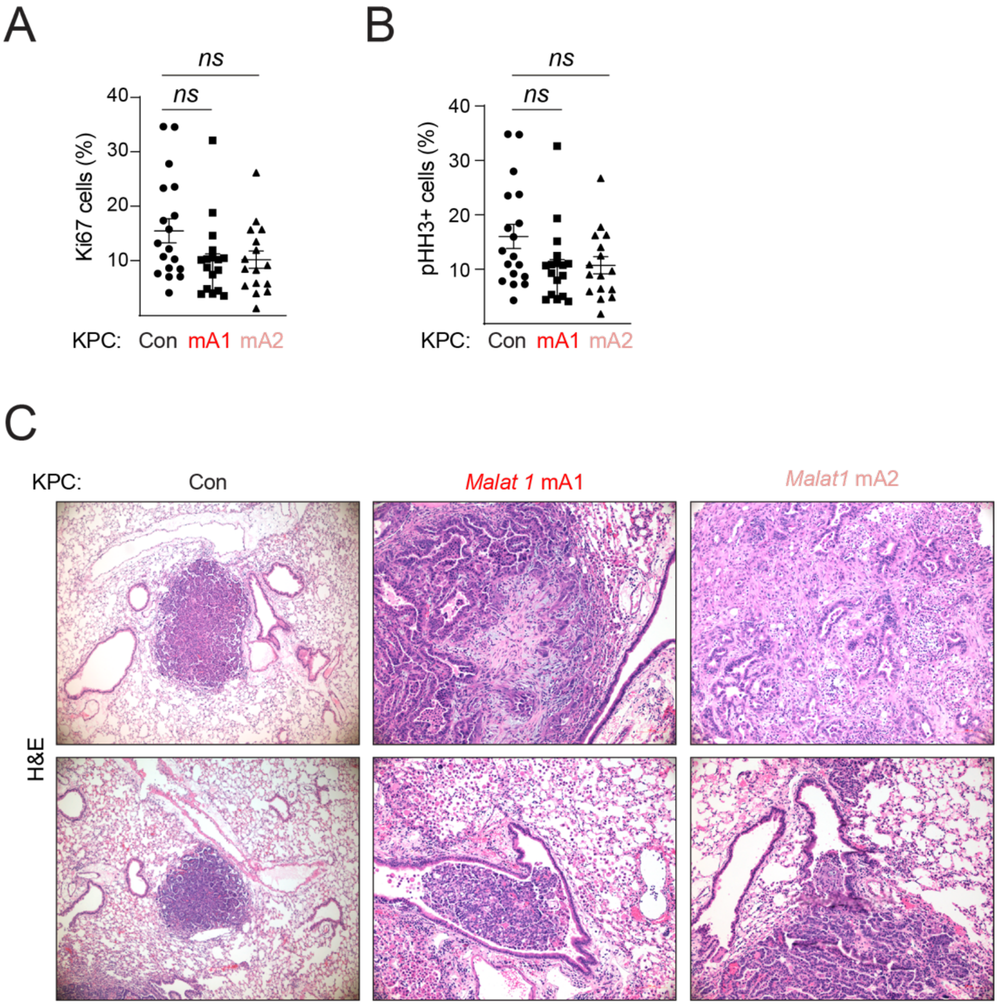


**<u>Figure S2.</u> *Malat1* overexpression promotes tumor progression without affecting cellular proliferation.** (A) Quantification of the proliferation marker Ki67 levels in lung sections from KPC animals, each symbol represents an individual tumor, unpaired t-test; (B) Quantification of the mitotic marker pHH3 in lungs sections from KPC animals, each symbol represents an individual tumor, unpaired t-test; (C) H&E staining of Con- mA1- and mA2- expressing KPC tumors with features of aggressive disease as shown by an extensive stroma infiltration (*Top*) and invasive tumors (*Bottom*) in mA1/2 KPC animals.

### Martinez-Terroba et al., Figure S3

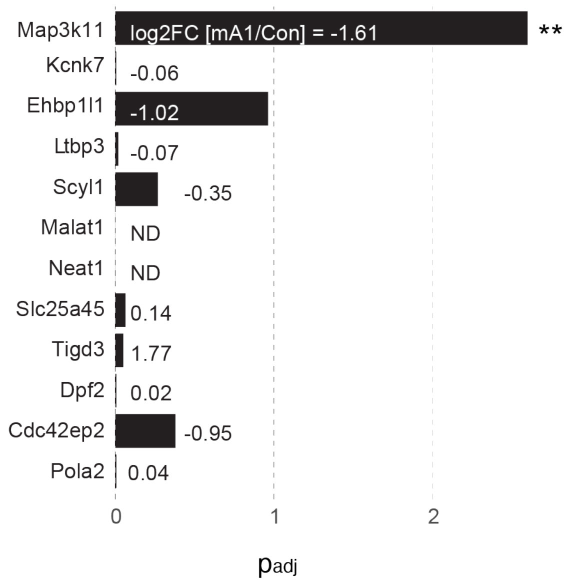


**<u>Figure S3.</u> *Malat1* activation affects global transcription *in trans*.** *Malat1* overexpression does not significantly affect the expression of neighboring genes with the exception of the significant downregulation of the oncogene, Mapk311. Numbers indicate log2 fold change in the expression of *Malat1* neighboring genes in mA1 compared to Con samples. ND, not determined, the levels of Neat1 and *Malat1* cannot be accurately determined due to their lack of polyA tails.

### Martinez-Terroba et al., Figure S4

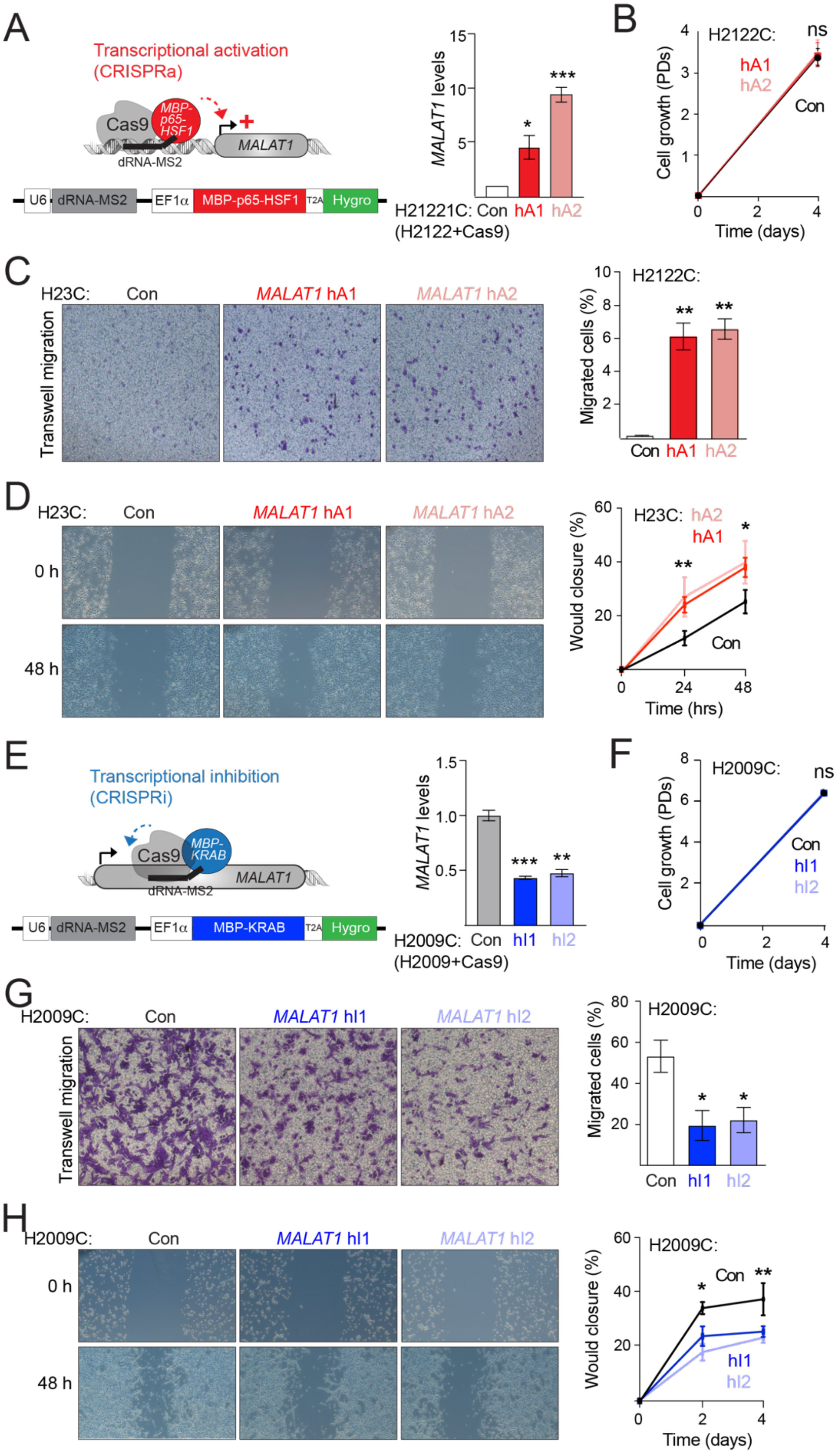

**<u>Figure S4.</u> *Malat1* overexpression is necessary and sufficient to promote cellular migration in human LUAD cell lines.** (A) *Top left*, schematic for the development of CRISPR-based tools for *Malat1* transcriptional activation (CRISPRa). *Bottom left*, Schematic of *in vitro* Lenti-SAM-Hygro construct. *Right*, RT-qPCR for *Malat1* RNA levels in H2122C cells in the absence (Con) and presence of *Malat1* activation (hA1/2); (B) Growth analysis of cells in (A); (C) *Left*, representative images of Boyden chamber transwell migration assay of indicated cells, scored in Fig. 4I. *Right*, quantification of transwell migration by Boyden chamber assay of H2122 cells; (D) Representative images and quantification of wound-healing scratch assay of cells in (C); (E) *Top left*, schematic for the development of tools for *Malat1* transcriptional inactivation (CRISPRi). *Bottom left*, schematic of *in vitro* Lenti-KRAB-Hygro construct. *Right*, RT-qPCR for *Malat1* RNA levels in the H2009C cells in the absence (Con) and presence of *Malat1* inactivation (hI1/2); (F) Growth analysis of cells in (E); (G) Representative images and analysis of Boyden chamber transwell migration assay of H2009C cells in (E); (H) Representative images and quantification of wound-healing scratch assay of cells in (E). Data show mean ± SEM (n = 3, biological replicates).

Martinez-Terroba et al., Figure S5

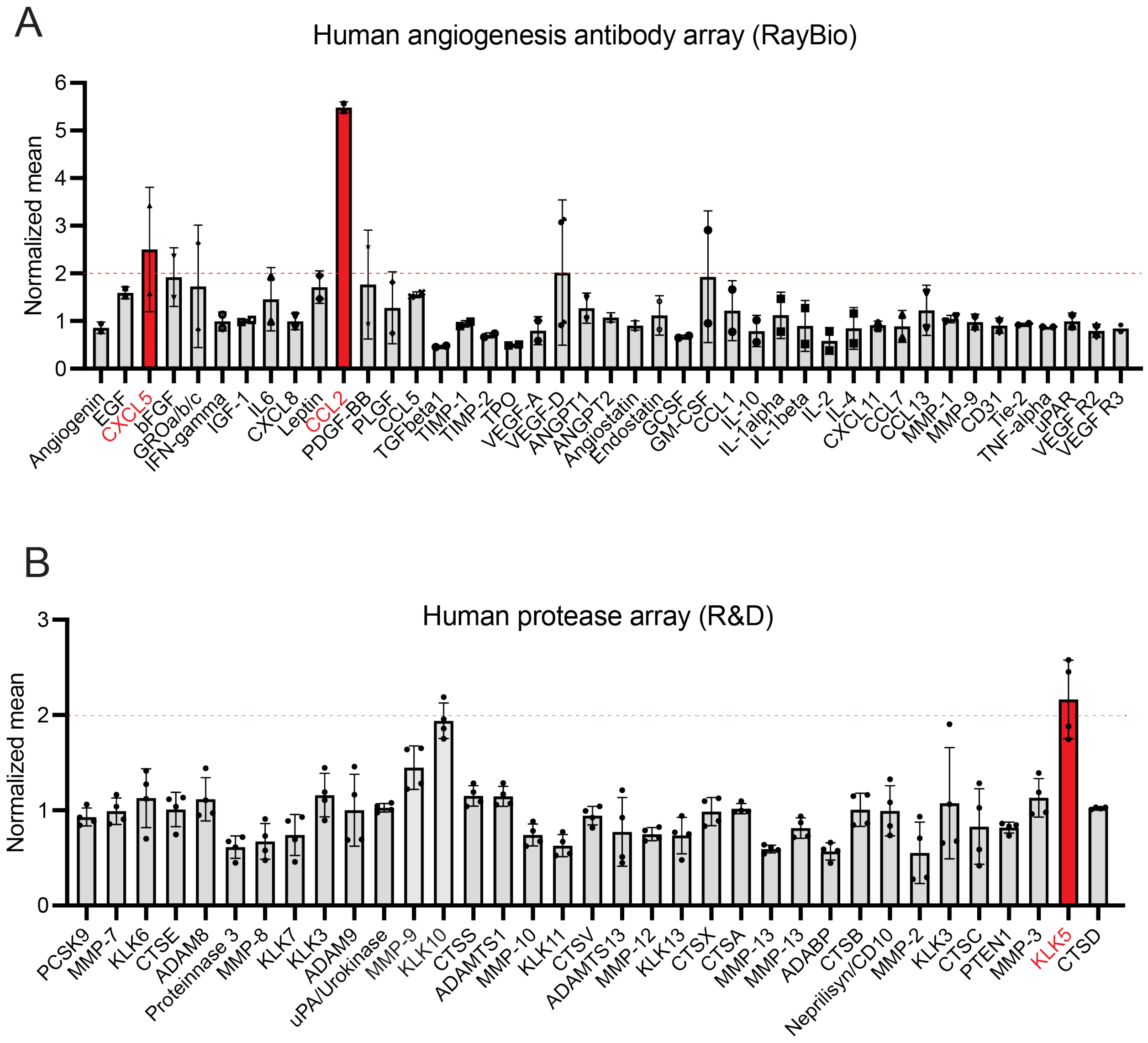


**<u>Figure S5.</u> Secretome changes resulting from *Malat1* overexpression.** (A-B) Quantification of human angiogenesis (A) and human proteases (B) antibody arrays incubated with conditioned media (CM) from Con or hA1 H23C cells. Con and hA1-incubated membranes were developed at the same time and normalized to internal controls. Densitometric analysis was used to evaluate differences in signal between Con or hA1-expressing cells.

Martinez-Terroba et al., Figure S6

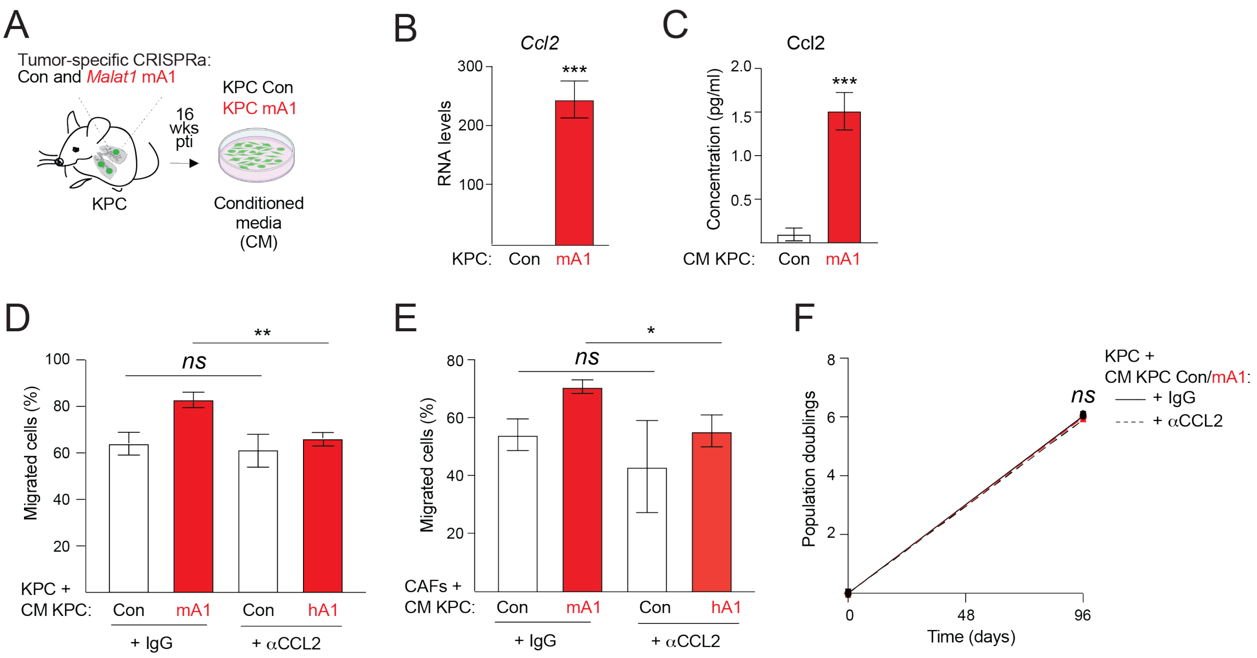


**<u>Figure S6.</u> The inflammatory cytokine Ccl2 mediates the paracrine effects of *Malat1* overexpression by enhancing migration of tumor and stromal cells.** (A) Schematic of the isolation of Con and mA1 KPC cell lines from KPC tumors; (B) *Ccl2* RNA levels in indicated KPC cells, (C) Ccl2 protein levels detected by ELISA in CM from indicated KPC cells; (D) Quantification of transwell migration assay of KPCs incubated with CM from indicated KPC cells in the absence (IgG) or presence of Ccl2-neutralizing antibody (αCcl2); (E) Quantification of transwell migration assay of CAFs incubated with CM from indicated KPC cells in the absence (IgG) or presence of Ccl2-neutralizing antibody (αCcl2); (F) Growth analysis of cells in (D).
